# Epigenetic neural glioblastoma enhances synaptic integration and predicts therapeutic vulnerability

**DOI:** 10.1101/2023.08.04.552017

**Authors:** Richard Drexler, Robin Khatri, Thomas Sauvigny, Malte Mohme, Cecile L. Maire, Alice Ryba, Yahya Zghaibeh, Lasse Dührsen, Amanda Salviano-Silva, Katrin Lamszus, Manfred Westphal, Jens Gempt, Annika K. Wefers, Julia Neumann, Helena Bode, Fabian Hausmann, Tobias B. Huber, Stefan Bonn, Kerstin Jütten, Daniel Delev, Katharina J. Weber, Patrick N. Harter, Julia Onken, Peter Vajkoczy, David Capper, Benedikt Wiestler, Michael Weller, Berend Snijder, Alicia Buck, Tobias Weiss, Michael B. Keough, Lijun Ni, Michelle Monje, Dana Silverbush, Volker Hovestadt, Mario L. Suvà, Saritha Krishna, Shawn L. Hervey-Jumper, Ulrich Schüller, Dieter H. Heiland, Sonja Hänzelmann, Franz L. Ricklefs

## Abstract

Neural-tumor interactions drive glioma growth as evidenced in preclinical models, but clinical validation is nascent. We present an epigenetically defined neural signature of glioblastoma that independently affects patients’ survival. We use reference signatures of neural cells to deconvolve tumor DNA and classify samples into low- or high-neural tumors. High-neural glioblastomas exhibit hypomethylated CpG sites and upregulation of genes associated with synaptic integration. Single-cell transcriptomic analysis reveals high abundance of stem cell-like malignant cells classified as oligodendrocyte precursor and neural precursor cell-like in high-neural glioblastoma. High-neural glioblastoma cells engender neuron-to-glioma synapse formation *in vitro* and *in vivo* and show an unfavorable survival after xenografting. In patients, a high-neural signature associates with decreased survival as well as increased functional connectivity and can be detected via DNA analytes and brain-derived neurotrophic factor in plasma. Our study presents an epigenetically defined malignant neural signature in high-grade gliomas that is prognostically relevant.

## INTRODUCTION

The importance of the nervous system as a key regulator of primary brain and metastatic tumors has been repeatedly highlighted but has not yet been translated into a therapeutically relevant setting^1^. The presence of neural-cancer interactions is a contributing factor in tumorigenesis and progression^1–3^. Particularly in gliomas, studies have demonstrated that the formation of malignant neuron-to-glioma networks is critical for cancer progression, and have identified crucial mechanisms such as paracrine signaling via neuroligin-3 (*NLGN-3*) or brain-derived neurotrophic factor (*BDNF*) and glutamatergic synapses driven by neuronal activity^3–6^. Additionally, glioma cells remodel neuronal circuits and are able to increase neuronal hyperexcitability^3, 7–10^. Therefore, targeting bidirectional neural-to-cancer interactions may be a promising therapeutic approach in poor prognosis gliomas, such as isocitrate dehydrogenase (IDH)-wildtype glioblastoma and H3 K27-altered diffuse midline glioma (DMG)^11, 12^.

Despite the increasing appreciation of the importance of neuroscience in understanding brain tumors, the targetable disruption of neuron-to-cancer synaptic communication in glioma was initially limited to preclinical models. Further insight into molecular mechanisms of neuron-to-glioma interactions identified connected and unconnected glioblastoma cells that form two distinct cell states and differ in their gene signatures as well as functions within neuron-to-glioma networks^13^. In addition, upregulation of neural signaling programs that promote brain tumor invasiveness at the time of recurrence has been demonstrated^14^. Recently, glioblastomas exhibiting high functional connectivity have been shown to be associated with poorer survival, and thrombospondin-1 (*TSP-1*)-expressing glioma cells have been identified as a key cell population for promoting neuron-to-glioma interaction^10^. Moreover, callosal projection neurons were shown to promote glioma progression and widespread infiltration underpinning the high importance of the central nervous system as a critical regulator and potential therapeutic target^15^.

High-grade gliomas are diffuse infiltrating tumors with a cellular composition consisting of both malignant and non-malignant cells which could be addressed by epigenetic bulk DNA analysis since it provides the possibility to decipher the underlying cellular composition. We hence hypothesized that by using brain tumor-related epigenetic signatures, we might decipher the epigenetic signature of IDH-wildtype high-grade gliomas and proposed that certain epigenetic subclasses may be more likely to be integrated into neuron-to-glioma networks and that their stratification may be clinically relevant. To address these hypotheses, we determined the tumoral neural signature by using a neural reference to screen bulk CNS tumors and stratified glioblastoma samples into low- and high-neural subgroups. These two distinct neural subgroups of glioblastoma were further molecularly, functionally, and clinically characterized by DNA methylation, spatial transcriptomics, single-cell deconvolution, proteomics, and imaging-based functional connectivity in human as well as *in vitro* and *in vivo* experiments.

We demonstrate that high-neural glioblastomas exhibit a synaptogenic profile and have an oligodendrocyte-precursor cell (OPC) and neuronal progenitor cell (NPC)-like character with a malignant stem cell-like state. High-neural glioblastomas show increased functional connectivity and neuron-to-glioma synapse formation *in vivo* and *in vitro*. Stratification of patients into low- and high-neural tumors proves to be an independent prognostic factor for survival in glioblastoma as well as DMG which highlights the clinical relevance of the here presented epigenetic neural signature.

## RESULTS

To address the aforementioned hypotheses, we applied the epigenetic neural signature of Moss et al^16^ to estimate cellular composition (Fig. 1a) of a combined dataset of epigenetically profiled central nervous system (CNS) tumors of Capper et al.^17^ and our institutional cohort (“clinical cohort”) (Supplementary Fig. 1). Using this combined dataset, IDH-wildtype glioblastoma samples (n=1058) were selected and dichotomized for defining a cut-off separating low- and high-neural tumors (cut-off 0.41, Supplementary Fig. 2). This cut-off was applied to 363 glioblastoma patients from our clinical cohort who received surgical treatment followed by standard-of-care combined chemo-radiotherapy. Survival analysis revealed a significantly shorter overall (OS) (p < 0.0001, median OS 14.2 versus 21.2 months, Fig. 1b) and progression-free survival (PFS) (p = 0.02, median PFS 6.2 versus 10.0 months, Fig. 1c) for patients with a high-neural glioblastoma (Supplementary Table 1). This finding was replicated in an external cohort with 187 patients from the TCGA-GBM database^18^who received adjuvant combined chemo-radiotherapy (p < 0.01, median OS 12.0 versus 17.1 months, Fig. 1d). Additionally, the neural classification was identified as an independent prognostic factor for OS (OR; 95% CI: 1.96; 1.45-2.64, p < 0.01, Fig. 1e) and PFS (OR; 95% CI: 1.51; 1.13-2.02, p < 0.01, Fig. 1f) next to established factors such as extent of resection (EOR), and O6-methylguanine-DNA-methyltransferase (*MGMT)* promoter methylation status (Supplementary Tables 2 and 3). Other infiltrating brain tumor cell types of the lymphoid or myeloid lineage did not show an association with patient survival (Supplementary Fig. 3).

**Figure 1:**
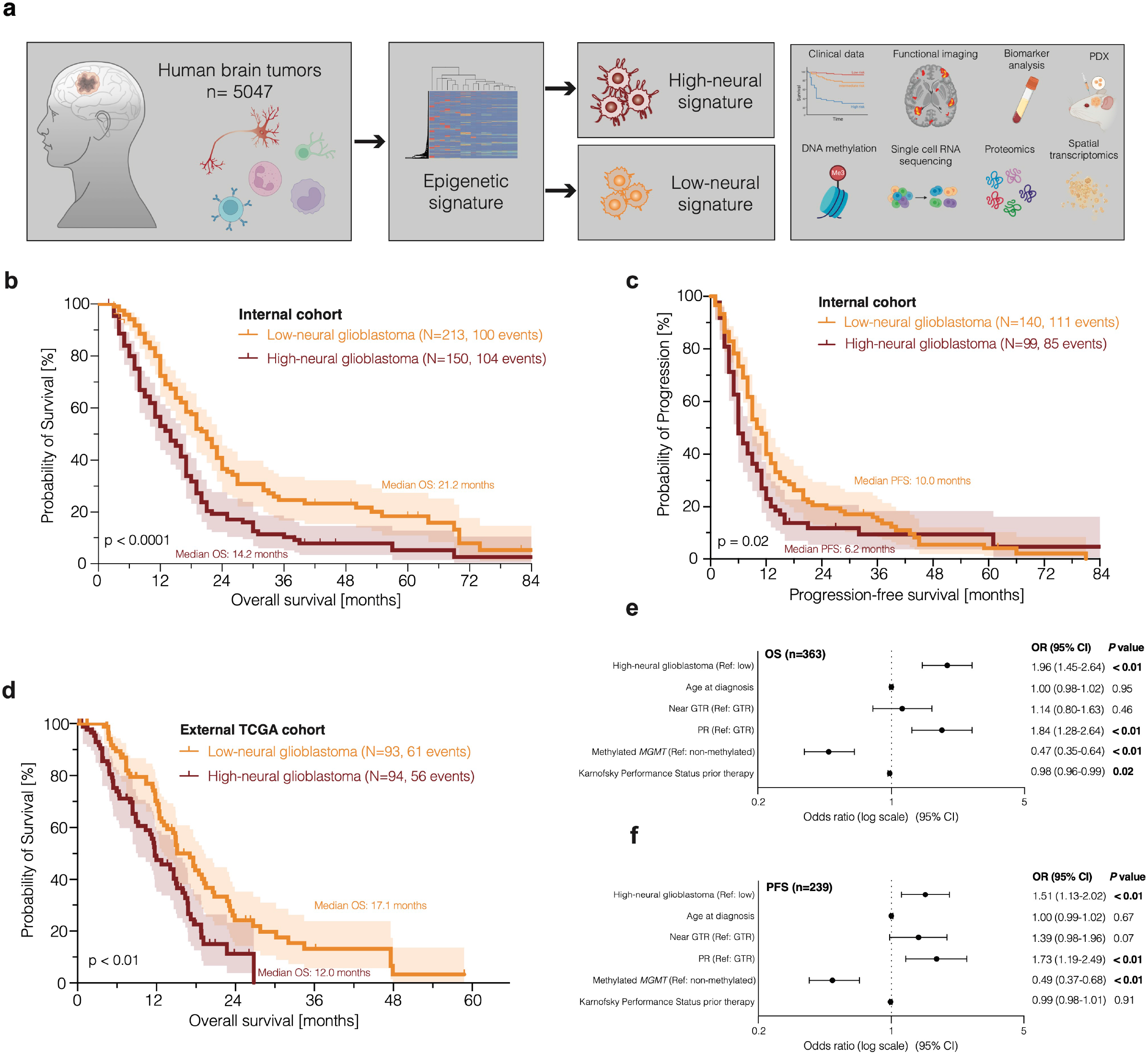
**a.) Schematic of study workflow.** In human subjects (n=5047) diagnosed with a central nervous system tumor we performed deconvolution using DNA methylation arrays (850k or 450k) for determining the neural signature. *IDH*-wildtype glioblastomas and *H3 K27*-altered diffuse midline gliomas were stratified into subgroups with a low- or high-neural signature for further analyses. **b.) – f.) Survival analysis of glioblastoma patients treated by radiochemotherapy after surgery with a low- and high-neural signature.** **b.)** Overall survival of 363 glioblastoma patients of the internal clinical cohort. **c.)** Progression-free survival of 226 glioblastoma patients of the internal clinical cohort. **d.)** Overall survival of 187 glioblastoma patients of the TCGA-GBM cohort. **e.) - f.)** Forest plots illustrating multivariate analysis of glioblastoma patients from the internal clinical cohort. *GTR: gross total resection, PR: partial resection, MGMT: O6-methylguanine-DNA-methyltransferase, OR: odds ratio, CI: confidence interval, TCGA: The Cancer Genome Atlas*.

### High-neural glioblastoma exhibits a synaptogenic and OPC-/NPC-like character

To further understand the survival difference and to demonstrate validity of the neural signature as a prognostic marker, we applied the “invasivity signature” by Venkataramani et al.^13^ which describes 172 genes associated with neural features, migration, and invasion (Extended Data 1) to the DNA methylation data of our clinical cohort. High-neural tumors were hypomethylated at CpG sites within gene loci of the invasivity signature (Fig. 2a). Additionally, two gene sets that are either associated with neuron-to-glioma synapse formation^11^ (“neuronal signature genes”, Extended Data 2) or relevant to trans-synaptic signaling^19^ (“trans-synaptic signaling genes”, Extended Data 3) were hypomethylated in high-neural glioblastomas (Supplementary Fig. 4a). Tumor DNA purity correlated with the neural signature, ruling out the possibility of sample contamination by non-malignant neural cells (p < 0.01, R^2^ = 0.38 Supplementary Fig. 4b).

**Figure 2:**
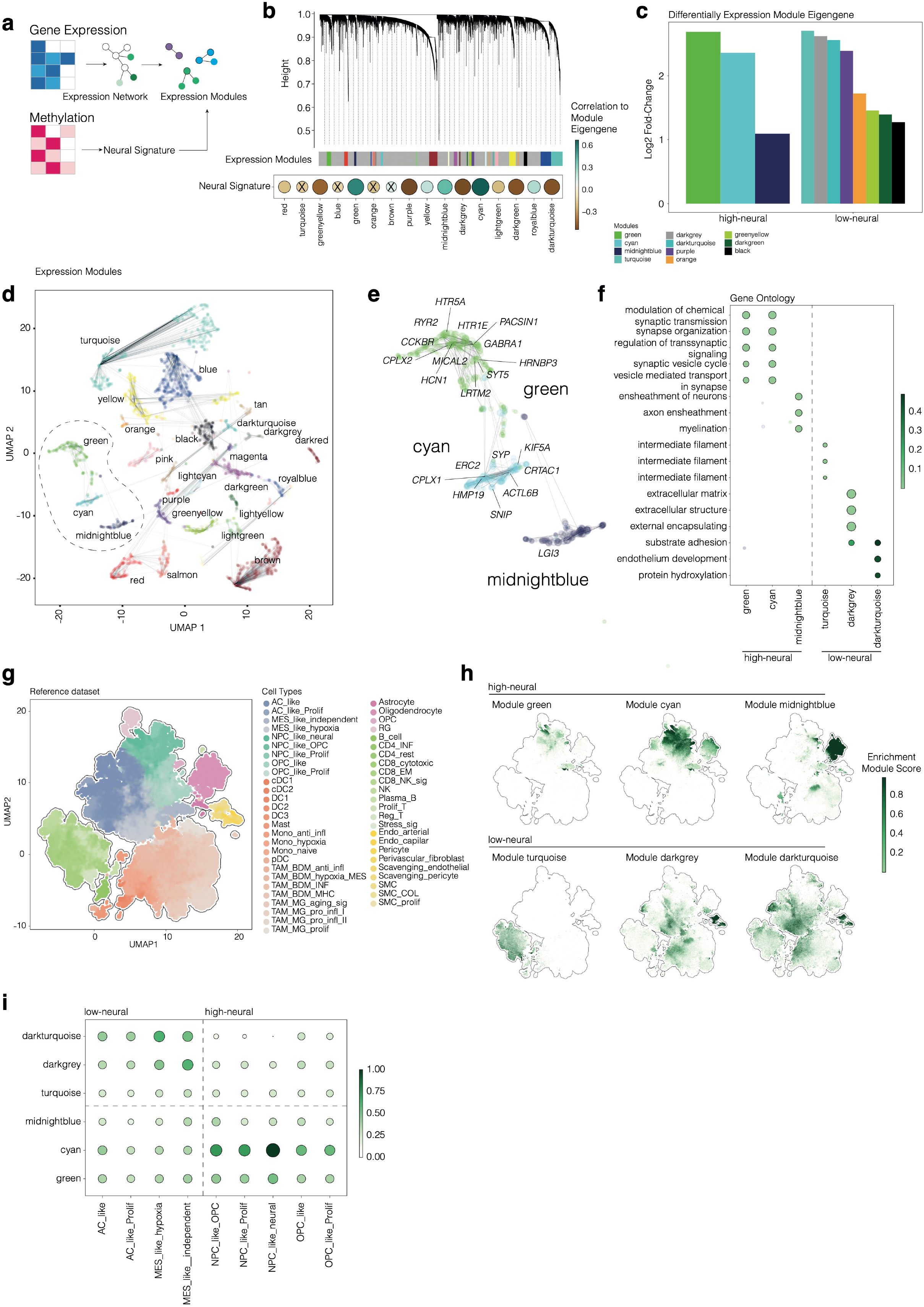
Integrated epigenetic and transcriptomic analysis reveals synaptic functions and a malignant neural precursor cell-like and oligodendrocyte precursor cell-like character in high-neural glioblastoma. **a.)** Illustration of the workflow to integrate epigenetic and transcriptional data. Gene co-regulation networks are correlated to the epigenetic deconvolution signature. **b.)** Hierarchical dendrogram of the gene expression modules derived from the weighted correlation network analysis. On the bottom, Pearson correlation dotplot of the neural signature with gene expression models. Size and color indicate the correlation coefficient, non-significant correlation is marked. **c.)** Barplot of the differentially gene expression of module eigengenes (Log2 fold change) in low- and high neural glioblastoma (cut-off 0.41). **d.)** Dimensional reduction (UMAP) of the gene expression modules (named by colors) and **e.)** a detailed visualization of the modules: green, cyan and midnightblue (significantly associated with high-neural tumors). **f.)** Gene ontology analysis of gene expression modules in low- and high-neural tumors. **g.)** UMAP dimensional reduction of the GBMap reference dataset. Colors indicate the different cell types. **h.)** Module eigengene expression of low- and high-neural glioblastoma in the GBMap reference dataset. **i.)** Gene expression enrichment of low- and high-neural associated module eigengenes across glioblastoma cell states.

Next, we used an integrative analysis of both epigenetic and transcriptomic datasets of glioblastoma samples (TCGA). Applying weighted correlation network analysis (WGCNA), we identified three expression modules significantly correlated with the epigenetic status of high-neural glioblastoma (Fig. 2a). Module green (R^2^=0.55 p=3.5 x 10^-6^), Module cyan (R^2^=0.67 p<2.2 x 10^-22^) and Module midnightblue (R^2^=0.41 p=9.3 x 10^-5^) (Fig. 2b-c). Gene ontology analysis revealed that these modules were associated with synaptic functions (*GRIN3A*, *SYT4*, *SNAP25*), regulating the expression of genes involved in neuronal differentiation (*NEUROD2*) and calcium-dependent cell adhesion (*CDH22*, *CNTNAP5* and *CNTN3*) (Fig. 2d-f). When projecting the module eigengene signatures onto a single cell dataset, malignant neural precursor cells (NPC)-like and oligodendrocyte precursor cell (OPC)-like (module green and cyan p<0.01) as well as non-malignant oligodendrocytes (module midnightblue p<0.01), revealed significant enrichment of the corresponding expression modules (Fig. 2g-i).

This pro-synaptogenesis signature of high-neural glioblastoma based on epigenetic and transcriptomic data was further validated in the tumor proteome by mass spectrometry analysis of 28 glioblastoma samples (low-neural: n = 18, high-neural: n = 10) (Supplementary Fig. 4c-h). High-neural glioblastoma exhibited increased proteins connected to synaptic transmission and vesicle-mediated transsynaptic signaling (Supplementary Fig. 4f). As previously seen in the spatial transcriptomic analysis, an OPC- and NPC-like character was evident in the high-neural glioblastoma cells after transfer to a single-cell data set (Supplementary Fig. 4g), as well as a malignant signature within these cells (Supplementary Fig. 4h).

To further investigate the spatial organization, we employed spatially resolved transcriptomics^20^ to samples which were epigenetically characterized as low- or high-neural glioblastoma (Fig. 3a-d). We observed a distinct spatial enrichment of the eigengene signatures from the green and cyan modules along with significant spatial correlation with the spatial OPC and neuronal development niches in high-neural tumors. Conversely, we confirmed the increased enrichment of low-neural glioblastoma expression modules in close relation to the necrotic core (Fig. 3a-b). These modules also spatially correlate with inflammation and metabolic alterations (Fig. 3c). To dissect the differences in cellular hierarchies and proximity between low- and high-neural glioblastoma, we computed cellular neighborhood graphs derived from single-cell deconvolution^21^ from samples which were epigenetically defined as low- and high-neural glioblastoma. Our observations indicated that the overall architecture of the tumors maintained similar (Fig. 3a-b), however, a more intricate interface between NPC/OPC-like cells and the non-malignant neuronal environment was evident only within high-neural glioblastoma (Fig. 3c-d).

**Figure 3:**
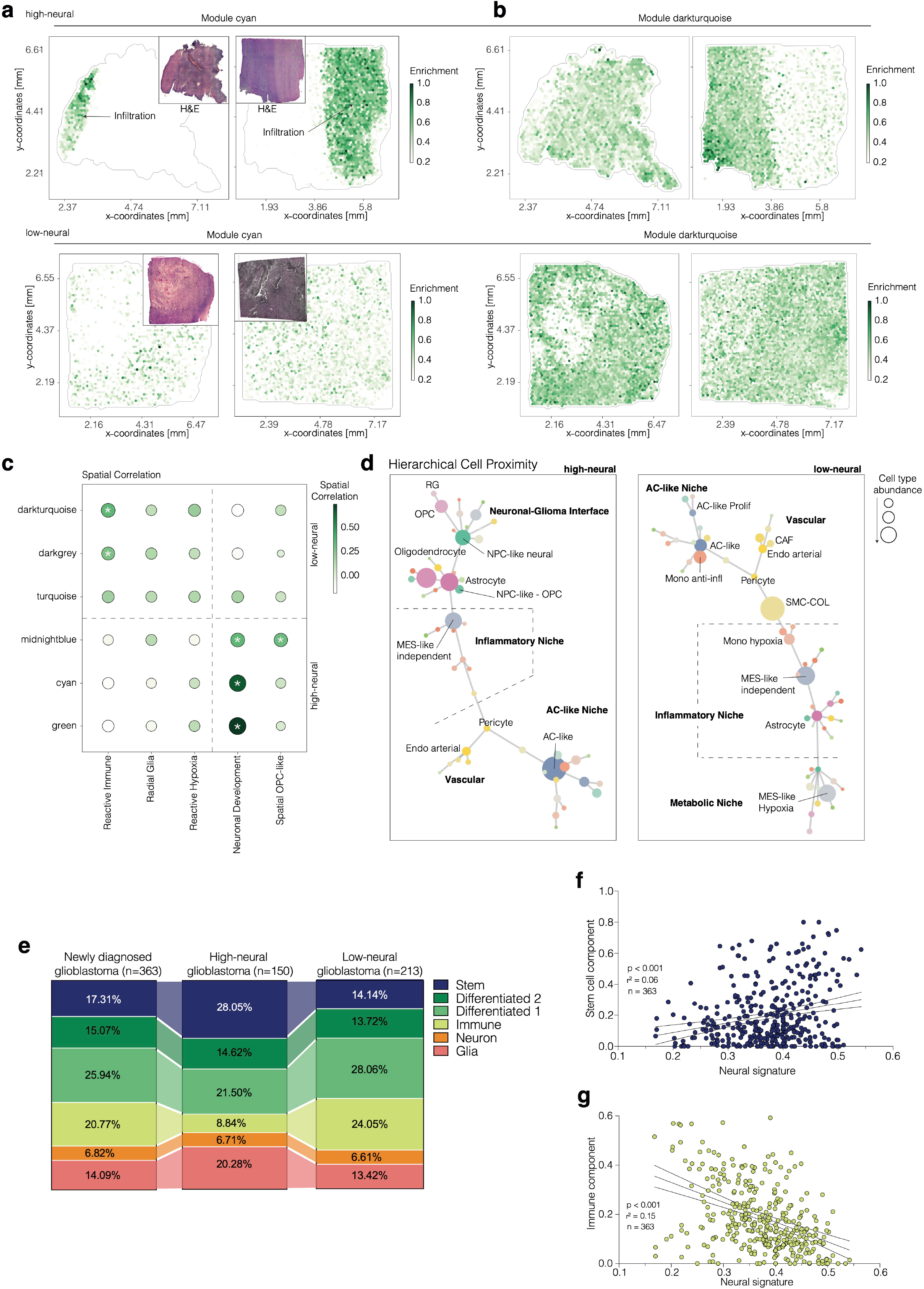
Spatial transcriptomic and single-cell deconvolution analysis of low- and high-neural glioblastoma samples. **a.) – b.)** Spatial transcriptomic surface plots of samples epigenetically defined as high (upper panel) and low (bottom panel) neural tumors. The colors indicate the spatial expression pattern of the module eigengenes. Representative H&E stainings are shown in the upper corner. **c.)** Spatial correlation analysis of low- and high-neural associated module eigengenes with the spatial transcriptomic niches. **d.)** Hierarchical cell architecture of high (left)- and low (right)-neural glioblastoma. Connections of the graph represent the spatial proximity of cell types and the size of the dots indicate the total cell type abundance and mapped the three upregulated modules to the infiltrative tumor zone. **e.)** Comparison of abundance of cell states analyzed by reference-free deconvolution between newly diagnosed, high-neural, and low-neural glioblastomas. **f.)** Stem cell-like state significantly correlated with an increase of the neural signature in glioblastoma samples. **g.)** An anticorrelation was seen between the abundance of the immune compartment and the neural signature.

### Analysis of the cell composition reveals an enriched stem cell-like state in high-neural glioblastoma

Brain tumor cells with a high neural state exhibit multiple neural features associated with neurodevelopmental programs^1^. We used a non-reference-based multi-dimensional single-cell deconvolution algorithm (see Methods) to further investigate the developmental status of our low- and high-neural glioblastoma samples. Here, a higher stem/progenitor cell-like component in the high-neural glioblastoma was observed (28.05%) compared to all newly diagnosed glioblastoma (17.31%) and low-neural glioblastoma (14.14%) (Fig. 3e). In contrast, the immune compartment was lower in high-neural glioblastoma (8.84% versus 20.77% versus 24.05%, Fig. 3e). We further determined a significant correlation of the neural signature with the stem cell component (p < 0.001, R^2^ = 0.06, Fig. 3f) and a significantly lower immune cell component (p < 0.001, R^2^ = 0.15, Fig. 3g). Copy number variations (CNV) of all glioblastoma samples were computed using the Conumee R package 1.28.0^22^. Tumors with a high and low-neural signature showed no significant differences in copy number variation (CNV) (Supplementary Fig. 5), further increasing the relevance of epigenetic signatures.

### High-neural glioblastoma engenders increased neuron-to-glioma synaptogenesis and worse survival in patient-derived xenograft models

Most studies elucidating the biology of cancer neuroscience in high-grade glioma were performed in preclinical models. We therefore examined the translatability of our neural classification to cell cultures and patient-derived xenograft (PDX) models. To this end, we analyzed the neural signature in cell cultures obtained from fresh samples of 17 glioblastoma patients and observed a well-preserved neural signature in 82.3% of our cell cultures compared to the original tumor samples (Fig. 4a-b). Analysis of cellular components by single-cell deconvolution revealed that *in vitro* culturing of tumor cells excluded the immune component and decreased the glial component, while the neural component remained stable, further supporting the epigenetically imprinted neural signature of glioblastoma cells (Fig. 4c).

**Figure 4:**
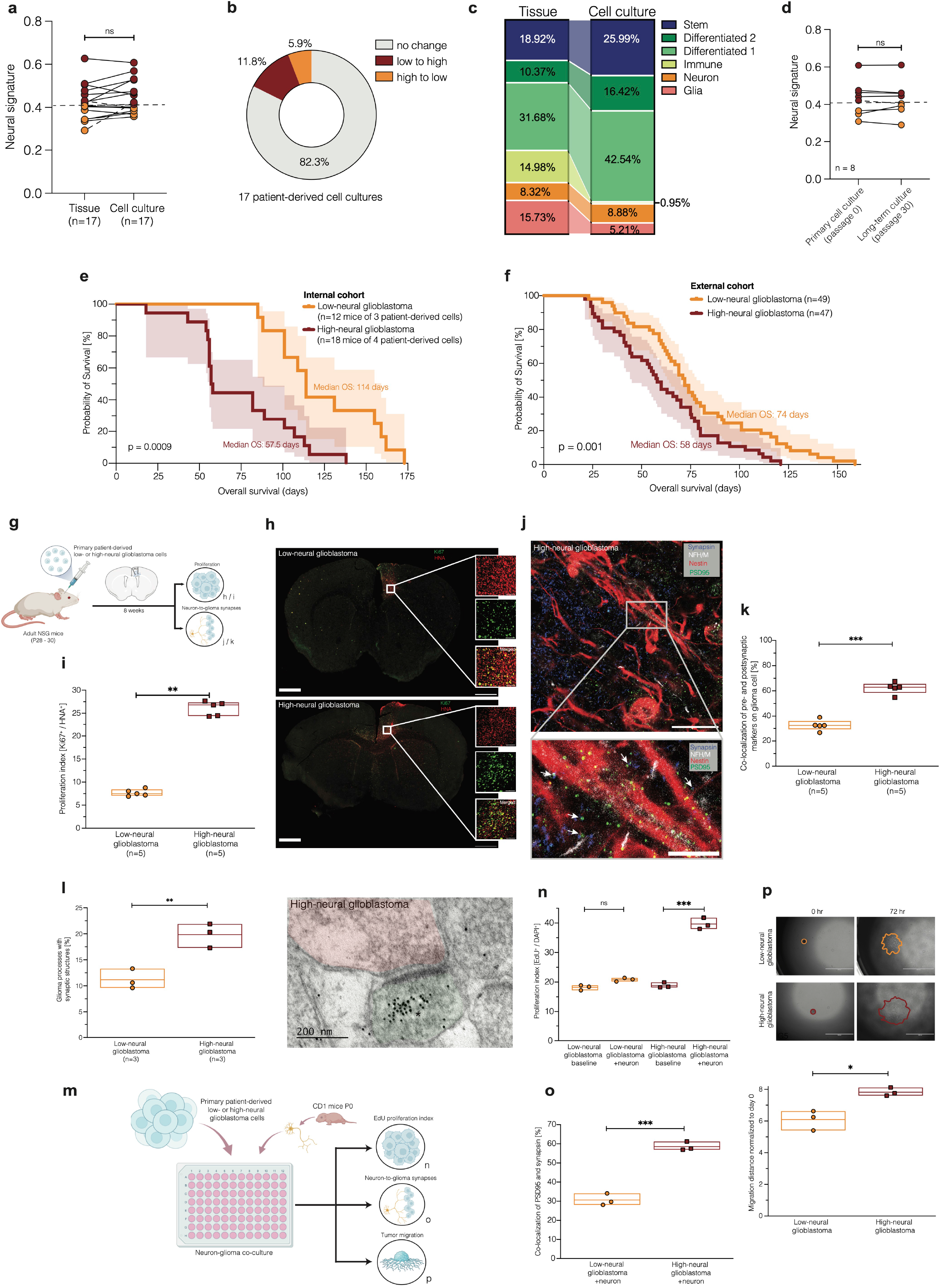
Neural classification is conserved in cell culture, correlates with survival *in vivo* and high-neural glioblastoma shows increased neuron-to-glioma synapses. **a.) – b.)** Comparison of neural signature between patient’s tumor tissue and cell culture in 17 glioblastomas. **c.)** Cell composition analysis represents the abundance of cell states between tumor tissue and cell culture. **d.)** Stability of the epigenetic neural signature during long-term cell culturing. Data were obtained from a publicly available dataset (n =6, GSE181314) and in-house (n = 1). **e.) – f.)** Mice survival after xenografting of patient-derived low- and high-neural glioblastoma cells in **e.)** our internal cohort, and **f.)** two combined external cohorts. **g.)** Primary patient-derived low- and high-neural glioblastoma cell suspensions (n = 1 per group) were implanted into premotor cortex (M2) of adult NSG mice (n = 5 mice per group). Mice were perfused after 8 weeks of tumor growth and brains sectioned in the coronal plane for further immunofluorescence analyses. **h.)** Representative confocal images of tumor burden in low-neural (upper image) and high-neural glioblastoma (bottom image) xenografts. *Human nuclear antigen (HNA), red; Ki67, green. Scale bars: 1.000 μm (overview images) and 200 μm (magnified images)*. **i.)** Proliferation index (measured by total number of HNA^+^ cells co-labelled with Ki67 divided by the total number of HNA^+^ tumor cells counted across all areas quantified) in low- and high-neural glioblastoma bearing mice (n = 5 mice per group). *\*\*P < 0.01, two-tailed Student’s t-test*. **j.)** Representative confocal image of infiltrated whiter matter of high-neural glioblastoma xenograft. White box and arrowheads highlight magnified view of synaptic puncta colocalization. *Blue, synapsin-1 (presynaptic puncta); white, neurofilament heavy and medium (axon); red, nestin (glioma cell processes), green, PSD95 (postsynaptic puncta). Scale bars: 500 μm (upper image) and 250 μm (lower image)*. **k.)** Quantification of the co-localization of presynaptic and postsynaptic markers in low- (n = 22 regions, 5 mice) and high- (n = 21 regions, 5 mice) neural glioblastoma xenografts. *\*\*\*P < 0.001, two-tailed Student’s t-test*. **l.)** Electron microscopy of patient-derived red fluorescent protein (RFP)-labelled low- and high-neural glioblastoma cells xenografted into the mouse hippocampus. Quantification of neuron-to-glioma synaptic structures as a percentage of all visualized glioma cell processes (left plot) and representative electron microscopy image of neuron-to-glioma process in a high-neural glioblastoma xenograft (right image). Asterix denotes immuno-gold particle labelling of RFP. Postsynaptic density in RFP^+^ tumor cell (pseudo-colored green), synaptic cleft, and vesicles in presynaptic neuron (pseudo-colored red) identify synapses. *\*\*P < 0.01, two-tailed Student’s t-test. Scale bar: 200 nm*. **m.)** Primary patient-derived low- and high-neural glioblastoma cells were co-cultured with cortical neurons from CD1-mice at P0 and further analyzed for proliferation and neuron-to-glioma synapse formation. Additionally, 3D migration assay was performed using monocultures of both cell lines. **n.)** EdU proliferation index (measured by total number of DAPI^+^ cells co-labelled with EdU divided by the total number of DAPI^+^ tumor cells counted across all areas quantified) in low- and high-neural glioblastoma as monocultures and co-cultured with neurons. *\*\*\*P < 0.01, ns: P > 0.05, two-tailed Student’s t-test, n=3 biological replicates*. **o.)** Quantification of the co-localization of PSD95 (postsynaptic) and synapsin-1 (presynaptic) in low- and high-neural glioblastoma cells in co-cultures with neurons. *\*\*\*P < 0.01, ns: P > 0.05, two-tailed Student’s t-test, n=3 biological replicates*. **p.)** 3D migration assay analysis with representative images at time 0 h (left) and 72 h (right) of low- and high-neural glioblastoma cells as well as comparison of distance of migration 72 h after seeding. *\*P < 0.05, two-tailed Student’s t-test, n=3 biological replicates. Scale bars: 1.000 μm*.

In addition, the neural signature remained stable in long-term cultures (p > 0.05, Fig. 4d). Comparison of low- and high-neural glioblastoma in PDX mouse models of an internal cohort (n=30 mice of 7 patient-derived glioblastoma cell cultures, Fig. 4e) and two publicly available cohorts^23, 24^ (n=96 patient-derived glioblastoma cell cultures, Fig. 4f) showed a significantly shorter survival of mice bearing high-neural tumors (internal cohort: p = 0.0009, external cohort: p = 0.001). These findings are consistent with the recent report of shorter survival in mice bearing orthotopic high functional connectivity (HFC) xenografts compared to those bearing low functional connectivity (LFC) xenografts^10^. In our study, an increased tumor burden assessed by Ki67^+^/HNA^+^ proliferation index (p < 0.01, Fig. 4g-i) and increased co-localization of neuron-to-glioma synapse puncta (p < 0.01, Fig. 4j-k) were seen in high-neural glioblastoma after injection of primary patient-derived cells of both neural subgroups in immunodeficient mice (n=5 per subgroup). The increased formation of neuron-to-glioma synapses in high-neural glioblastoma was additionally proven using electron microscopy in red fluorescent protein (RFP)-labelled, patient-derived low- and high-neural xenografts (n = 3 per group, p = 0.008, Fig. 4l). In accordance with our *in vivo* experiments, an increased proliferation of high-neural glioblastoma cells but not low-neural glioblastoma cells was seen when co-cultured with neurons (p < 0.001, Fig. 4m-n). Furthermore, we found an increase of co-localization of synapse puncta in high-neural glioblastoma cells (p < 0.001, Fig 4o), supporting the previously mentioned findings after xenografting. Since neuronal activity has recently been shown to be a factor in widespread infiltration of glioblastoma cells^15^, we wondered if this was also a characteristic of the high-neural glioblastomas in our study. For this purpose, we performed a migration assay, in which a significantly wider migration of high-neural glioblastoma cells could be demonstrated (p < 0.05, Fig. 4p).

The translation of the neural signature into cell cultures and PDX models demonstrates the robustness of the epigenetically imprinted neural signature and indicates its distinct role within neuron-to-glioma networks of the high-neural glioblastoma subgroup.

### High neural glioblastoma shows increased tumor connectivity and remains spatiotemporally stable

Previous studies have reported a relationship between tumor connectivity and patient survival^10, 25^. Here, we measured functional tumor connectivity using magnetoencephalography (n = 38, Fig. 5a-b) and resting state functional magnetic resonance imaging (n = 44, Fig. 5c-e) in glioblastoma patients. Both modalities showed a significant association of higher connectivity with the high-neural subgroup (p < 0.01, Fig. 5a-e). These findings are consistent with a recent study of distinct cellular states in regions of HFC-glioblastoma^10^. Comparing the functional connectivity phenotype^10^ to our neural classification, we found high concordance between both classifications. Volumetric analysis showed significantly smaller volumes of contrast-enhancement (p = 0.03, Fig. 5f) in high-neural glioblastoma, but no association with fluid attenuated inversion recovery (FLAIR) signal abnormality volume (p = 0.18, Fig. 5g) and necrotic volume (p = 0.78, Fig. 5h).

**Figure 5:**
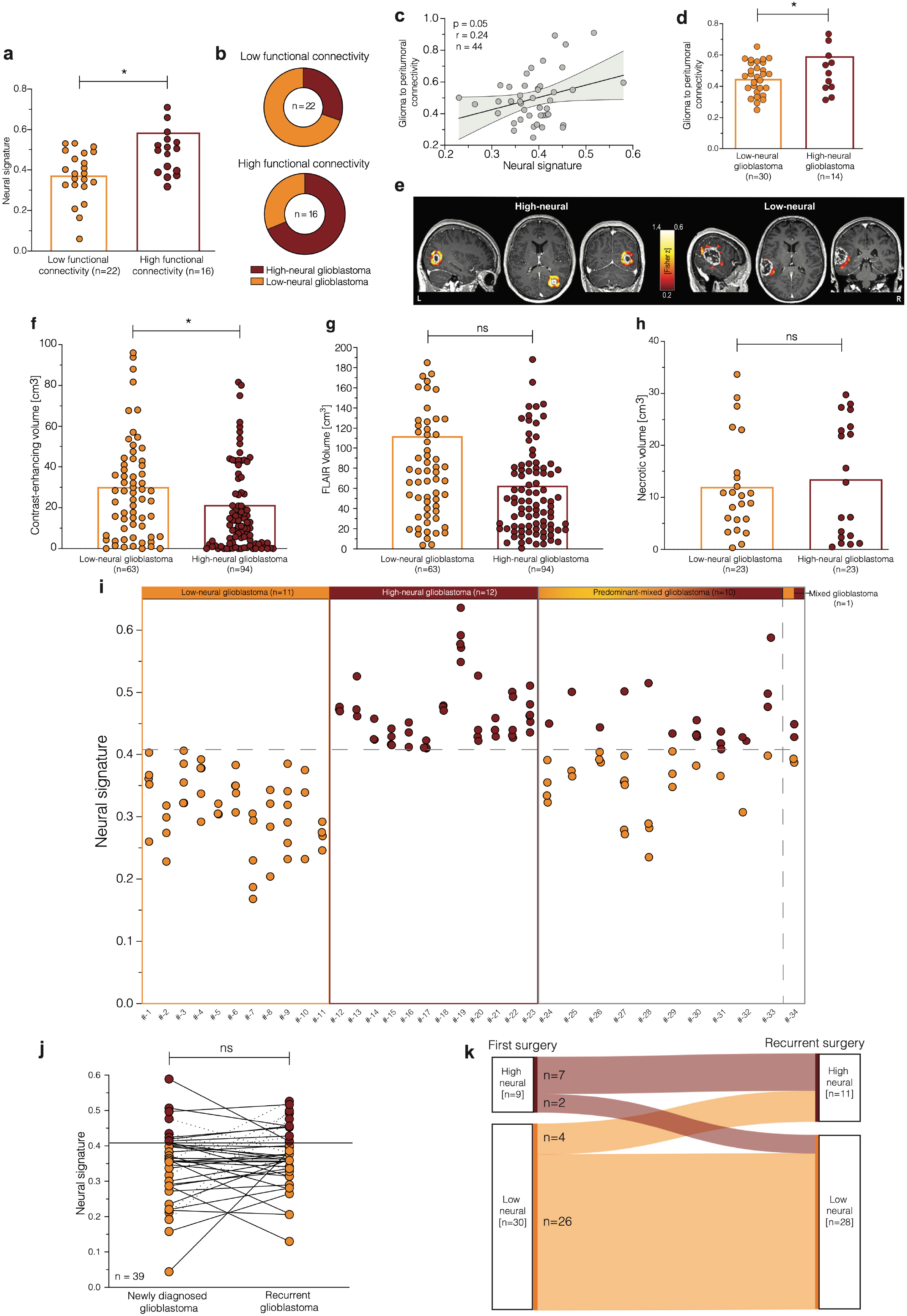
Association of the neural signature with functional connectivity and spatiotemporal stability in glioblastoma. **a.)** Neural signature in glioblastomas categorized into low (LFC) and high functional connectivity (HFC) as defined by magnetoencephalography. *\*P < 0.05, two-tailed Student’s t-test*. **b.)** Overlap between tumor samples classified to the functional connectivity by Krishna et al. and the epigenetic-based neural classification of our study. **c.)** Correlation of neural signature with degree of glioma to peritumoral connectivity as defined by resting state functional magnetic resonance imaging. **d.)** Correlation between functional connectivity as defined by resting state functional magnetic resonance imaging and low- and high-neural glioblastoma groups. *\*P < 0.05, two-tailed Student’s t-test*. **e.)** Two representative examples of patients with glioblastoma showing the ROI-to-voxel functional connectivity of the CE-enhancing area (ROI) to its 10mm peritumoral surrounding. Left image shows the peritumoral connectivity of patient with high-neural score (0.457) and mean functional connectivity to its peritumoral area of 0.837. In contrast, the right panel shows a patient with a low-neural score (0.347) and mean functional connectivity to its peritumoral area of 0.294. **f.) – h.)** Association of neural glioblastoma group with volume of **f.)** contrast-enhancement, **g.)** FLAIR, and **h.)** tumor necrosis measured by preoperative magnetic resonance imaging. *\*P < 0.05, ns: P > 0.05, two-tailed Student’s t-test*. **i.)** Analysis of intertumoral difference of neural signature within 34 newly diagnosed glioblastomas with spatial collection of 3 to 7 samples per tumor. 23 (67.6 %) of these tumors had a pure low- or high-neural signature in all individual biopsies with additional 10 (29.4 %) tumors being predominantly low or high. **j.)** Neural signature in 39 patients with matched tumor tissue obtained from surgery at first diagnosis and recurrence. *ns: P > 0.05, two-tailed Student’s t-test*. **k.)** Sankey plot illustrating a potential switch of the neural subgroup between first diagnosis and recurrence.

To address the topic of spatiotemporal heterogeneity, we analyzed spatially collected biopsies (3 to 7 samples of 34 patients, n = 143). Among them, 23 patients (67.6%) had a pure low- or high-neural signature, and a predominant signature was present in an additional 10 patients (29.4%) (Fig. 5i). To describe temporal stability, neural signatures were analyzed in 39 patients with matched tissue obtained from first and recurrence surgery (Fig. 5j-k). Here, 31 of 39 patients (79.5%) were categorized in the same neural subgroup at recurrence as at the time of diagnosis (Fig. 5k). Overall, the neural subgroup appeared to be spatiotemporally stable, in contrast to transcriptional states that change in a larger proportion of patients^14, 26^.

### Drug sensitivity analysis of neural glioblastoma cells

Glioblastoma patients routinely undergo combined radio-chemotherapy after surgical resection^27^. We therefore evaluated 27 different agents for their efficacy in the treatment of low- and high-neural glioblastoma cells (Supplementary Fig. 6a). We observed a trend for increased cleaved caspase 3 (Supplementary Fig. 6b) and reduced tumor cell size (Supplementary Fig. 6c) after treatment with lomustine (CCNU), JNJ10198400, and cyclosporine-treated high-neural glioblastoma cells, whereas talazoparib showed a trend for greater sensitivity in low-neural glioblastoma cells. However, none of these compounds reached statistical significance (Supplementary Fig. 6d). Therefore, we wondered about the prognostic impact of surgical resection in low- and high-neural glioblastoma since surgery is a cornerstone of glioblastoma therapy, and we previously demonstrated survival differences for other methylation-based glioblastoma subclasses^28^.

### Neural classification predicts benefit of resection in glioblastoma

Glioblastomas are epigenetically assigned to different subclasses with *receptor tyrosine kinase* (*RTK*) *I*, *RTK II* and mesenchymal (MES) being the most prominent in adult patients^29^. Here, *RTK I* and *RTK II* tumors showed a comparable neural signature while MES tumors had the lowest neural signature (Fig. 6a). Given the different neural signatures between methylation-based subclasses, we hypothesized that the neural signature might constitute a factor for determining benefit from different categories of extent of resection (EOR). In low-neural glioblastoma, a significant survival benefit of gross total resection (GTR) (100% CE resection) and near-GTR (≥90% CE resection) was observed compared with partial resection (PR; <90% CE resection) (p < 0.001, Fig. 6b). In contrast, the survival benefit of a near-GTR was not seen in high-neural glioblastoma (Fig. 6c). To further validate the differential benefit at distinct extents of resection in the two neural subgroups, we applied the current criteria of the Response Assessment in Neuro-Oncology (RANO) resection group^30^ to a subset of 174 glioblastomas from our clinical cohort.

**Figure 6:**
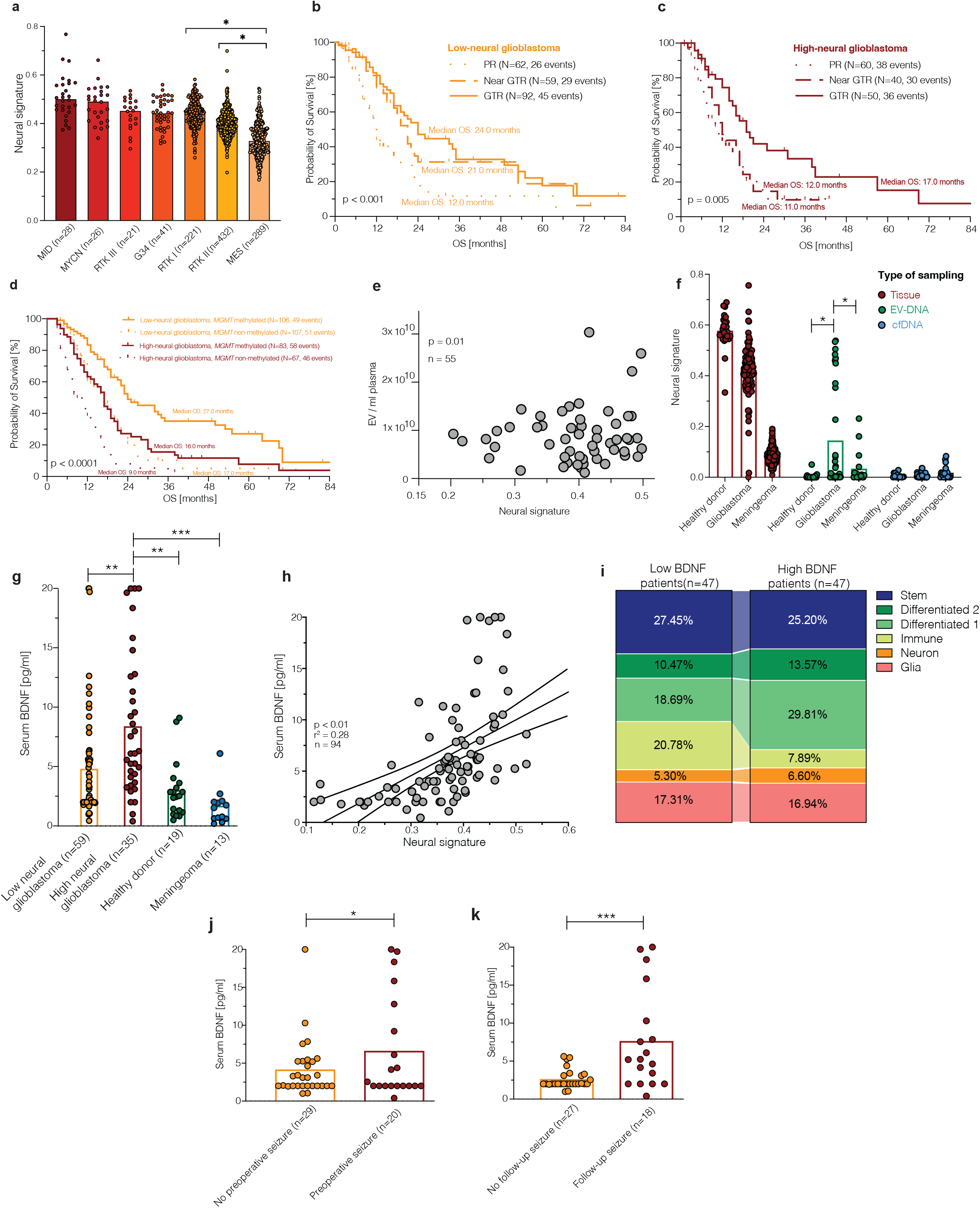
Neural classification predicts benefit of extent of resection and *MGMT* promoter methylation status and can be detected in serum of glioblastoma patients. **a.)** Neural signature in DNA methylation subclasses of newly diagnosed IDH-wildtype glioblastoma. **b.) – c.)** Survival outcome categorized after extent of resection in glioblastoma patients treated by radiochemotherapy with a **b.)** low- and **c.)** high-neural tumor. **d.)** Survival outcome categorized by *MGMT* promoter methylation status in glioblastoma patients treated by radiochemotherapy with a low- and high-neural tumor. **e.)** Correlation of neural signature and number of extracellular vesicles in patient serum at time of diagnosis. **f.)** Comparison of neural signature in healthy individuals, glioblastoma patients, and meningeoma patients between matched tumor tissue, extracellular vesicle-associated DNA in serum, and cell-free DNA in serum. **g.) – h.)** Immunoassay quantification of serum BDNF concentration of 94 glioblastoma patients and healthy donors as well as meningioma patients as control groups at time of diagnosis. *\*\*P < 0.01, ***P < 0.001, two-tailed Student’s t-test*. **i.)** Cell composition analysis in glioblastoma with low and high BDNF serum levels. **j.) – k.)** Seizure outcome of glioblastoma patients considering BDNF serum levels **j.)** at time of surgery, and **k.)** during follow-up. *\*P < 0.05, ***P < 0.001, two-tailed Student’s t-test*.

Here, again, it was found that the benefit of the category for extent of resection depends on the neural subgroup (Supplementary Fig. 7). While an extent of resection of category 3A (≤ 5cm CE) showed a significant survival benefit in patients with a low-neural glioblastoma compared to category 3B (≥ 5cm CE), this was not evident in the high-neural tumors (Supplementary Fig. 7). Consideration of the *MGMT* promoter showed a survival benefit of a methylated promoter in both subgroups, but a striking difference in low-neural glioblastoma with a median OS difference of 12.0 months depending on the *MGMT* promoter methylation status (p < 0.0001, Fig. 6d). Our combined survival data demonstrate that glioblastomas with a high-neural signature have an unfavorable survival prognosis, and a greater resection of contrast-enhancing tumor areas may be required to achieve a survival benefit in this distinct glioblastoma subclass.

### Serum biomarkers of high-neural glioblastoma

Further, we investigated whether preoperative assessment of the neural subgroup is feasible in blood. For this purpose, we determined the neural signature of circulating DNA analytes (extracellular vesicles (EV)-associated DNA and cell-free DNA (cfDNA)) in plasma of glioblastoma patients (Fig. 6e-f). Healthy individuals and meningioma patients were used as controls. Circulating EVs, a known blood-derived surrogate marker for tumor presence in glioblastoma^31^ and involved in neuron-to-glioma synchronization^32^, correlated with the neural signature of the distant glioblastoma (p < 0.01, n=55, Fig. 6e). Epigenetic profiling of EV-DNA in plasma showed a detectable neural signature that was not evident in cfDNA (Fig. 6f). Additionally, the neural signature in EV-DNA was significantly increased in glioblastoma compared to the two control groups (Fig. 6f). Apart from the detection of the neural signature in patient serum, brain-derived neurotrophic factor (BDNF) was repeatedly demonstrated as one factor for promoting neuronal activity-regulated glioma growth and reinforcing neuron-to-glioma interactions^4, 33^. We determined BDNF serum levels from 94 glioblastoma patients at time of diagnosis (Fig. 6g-h).

Patients with high-neural glioblastoma exhibited elevated BDNF serum levels compared to low-neural glioblastoma as well as meningioma (n=13) and healthy donors (n=19) (Fig. 6g). Glioblastomas with higher BDNF serum levels had a decreased immune cell signature, consistent with the low immune cell signature of high-neural tumor tissue samples (Fig. 6i).

Since neuronal activity increased BDNF release and current literature describes BDNF elevation in serum after provoked seizures for electroconvulsive therapy^34, 35^, we hypothesized that tumor-associated epilepsy may also promote BDNF release. Here, we observed a significant increase of BDNF levels in patients with epileptic seizures at time of diagnosis (p = 0.02, Fig. 6j) and during follow-up (p < 0.001, Fig. 6k).

These data suggest that EV-DNA and BDNF may serve as serum markers to stratify glioblastoma patients according to their neural subgroup and can be utilized for potential future targeted therapies.

### Epigenetic neural classification informs patients survival in H3 K27-altered diffuse midline glioma

Besides glioblastoma, the importance of neuronal activity in promoting tumor growth in DMG has been highlighted in previous studies^4, 5^. Therefore, we aimed to identify the neural signature in an additional cohort of H3 K27-altered DMG. The cohort consisted of patients from our institutional cohort (n=21), Chen et al.^36^ (n=24), and Sturm et al.^29^ (n=10). The neural signature was evenly distributed among tumors in the thalamus, pons, and medulla (Supplementary Fig. 8a). As previously observed for glioblastomas, CpG sites within the genes associated with invasivity, neuron-to-glioma synapse formation, and transsynaptic signaling were predominantly hypomethylated in high-neural DMG (Supplementary Fig. 8b). Additionally, cell state composition analysis showed a higher immune component in low neural tumors, whereas high-neural DMG samples were associated with stem and glial cell states (Supplementary Fig. 8c). Further, the malignant stem cell-like, OPC-like state was found to be correlated with synaptic gene expression in a single-cell RNAseq dataset by Venkatesh and colleagues (p = 0.01, r^2^ = 0.40, Supplementary Fig. 8d)^3^. Survival analysis of 72 pediatric and adolescent patients showed an unfavorable outcome for high-neural DMG (p < 0.01, Supplementary Fig. 8e). The survival difference between low- and high-neural DMG was significant when localized in the thalamus (p < 0.01, Supplementary Fig. 8f) but not in the pons (p = 0.08, Supplementary Fig. 8h) and medulla (p = 0.32, Supplementary Fig. 8g).

These results for patients with a DMG are consistent with the previous findings in glioblastoma and confirm the relevance of the neural signature in an additional type of IDH-wildtype high-grade glioma.

## DISCUSSION

In recent years, the bidirectional interaction between glioma cells and neural cells, with their ability to form synapses and integrate into neuronal circuits, has been identified as a major factor in oncogenesis and tumor progression^3, 4, 13, 37^. In this study, we identified an epigenetically defined malignant neural signature as a potential marker for neural-to-glioma interactions among glioblastoma and DMG and present the following findings: 1. A malignant neural signature is increased in glioblastoma and DMG, compared with non-malignant brain tumors. 2. High-neural glioblastoma confers an unfavorable survival in humans and mice, and in addition, the neural signature is associated with higher functional connectivity in glioblastoma patients. 3. High-neural glioblastoma shows an increased malignant stem cell and NPC/OPC-like character but decreased immune infiltration. 4. The neural signature remains robust in PDX mouse models and high-neural glioblastoma bearing mice show higher proliferation and migration as well as increased neuron-to-glioma synapses. 5. High-neural tumors benefit from a maximized resection. 6. The epigenetic neural signature can be detected in circulating EVs. 7. Elevated BDNF serum levels are present in high-neural glioblastoma and are associated with a higher rate of preoperative and therapy-refractory seizures. 8. The neural signature and its prognostic value can also be seen in DMG, an additional IDH-wildtype malignant glioma tumor type.

Gliomas encompass a variety of cellular components of the tumor microenvironment, and subgroups can be described according to distinct cellular states^38^. In addition, epigenome profiling and deconvolution were shown to characterize the microenvironment of glioma methylation subclasses^39, 40^. We further distinguished IDH-wildtype gliomas according to their epigenetic neural signature as a potential marker of neuron-to-glioma interactions. An increase in neural signature was found in glioblastoma and DMG, which reflects the findings of previous studies in preclinical models^3, 5^.

By multi-dimensional profiling, high-neural glioblastoma showed upregulation and hypomethylation of genes known to be associated with invasiveness and neuron-to-glioma synapse formation and signaling. It is well established that glioma growth occurs through paracrine signaling and glutamatergic synaptic input^3–6, 33^, and recently Venkataramani and colleagues subdivided glioblastoma cells into unconnected and connected cells with unique cell states, explaining brain infiltration through hijacking of neuronal mechanisms^13^. Spatial transcriptomic analysis revealed a malignant OPC/NPC-like character of high-neural glioblastoma cells consistent with the unconnected glioblastoma cells described by Venkataramani that hijack neuronal mechanisms and drive brain invasion. Additional cell state composition analysis profiled these high-neural tumors with a malignant stem cell-like character. Of note, the observed diploid oligodendrocyte transcriptomic module may represent a tumor cell population of primary near-diploid state as glioblastomas are karyotypically heterogeneous tumors, composed of many cellular populations^41^.

The clinical relevance of our findings is supported by the observation that patients suffering from high-neural glioblastoma or DMG had an unfavorable overall and progression-free survival. In addition, a greater extent of resection must be achieved to have prognostic improvement in high-neural glioblastoma, which may explain the results of our previous study examining the impact of DNA methylation subclasses^28^. Our findings are in line with a recent study by Krishna and colleagues which also showed poorer survival in patients with glioblastoma that exhibited high functional connectivity.^10^ Translating our signature to samples from Krishna et al. related an increased functional connectivity to a higher neural signature. The findings of this translational approach between both studies highlights *TSP-1*, a crucial driver of functional connectivity identified in Krishna’s study, as a potential therapeutic target. To further address the importance of tumor connectivity, we integrated glioblastoma patients who underwent preoperative resting state functional MRI and could also find an increased connectivity of high-neural glioblastomas to its peritumoral surrounding.

The synaptogenic character with increased functional connectivity of high-neural glioblastomas could be replicated with *in vivo* and *in vitro* experiments. Collectively, these data underscore the tremendous importance of the synaptic integration of gliomas into neuronal circuits and targeting these neuron-to-glioma networks appears to be a promising therapeutic approach.^1, 12^

One factor raising attention is *BDNF*, a neuronal activity-regulated neurotrophin, which has been found to promote glioma growth^4, 42^. Taylor et al. characterized BDNF as an enhancer of neural-glioma interactions and demonstrated therapeutic potential of interrupting BDNF-TrkB signaling in pediatric IDH-wildtype glioblastoma and DIPG^33^. Here, we found elevated serum BDNF levels in adult patients with a high-neural glioblastoma. Potential sources of elevated BDNF include neurons in a glioma-induced state of hyperexcitability^3^, given the known activity-regulation of BDNF expression and secretion^43–45^ or possibly from glioblastoma cells, as a subset of glioblastoma cells express and secrete BDNF^46^. Additionally, and consistent with findings in preclinical models, elevated serum BDNF levels were associated with a higher seizure frequency. The relationship between BDNF and seizure outcome fits with previously published data, as on the one hand BDNF regulates trafficking of *AMPAR* to the postsynaptic membrane of glioma^33^ and on the other hand an upregulation and hypomethylation of *AMPA* genes was found in the *RTK II* subclass, a highly epileptogenic glioblastoma subclass^47, 48^. Here, neuronal activity arising from glioma-to-neuron interactions during tumor growth or the onset of seizures seems to be a pivotal driver for BDNF release, as increased BDNF serum concentrations have already been shown after artificial induction of activity by electroconvulsive therapy.^34, 35^ Briefly summarized, these results identify a biomarker of high-neural glioblastoma, underline the importance of BDNF in glioma progression as well as tumor-related epilepsy, and highlight disruption of BDNF-TrkB signaling as a therapeutic target.

While this axis may represent a therapeutic target for high-neural glioblastoma, we further identified low-neural tumors as immune-enriched based on transcriptomic and cell state composition analysis. Consequently, one could hypothesize that two opposing glioblastoma subtypes appear to be differentiated here and will need to be pursued in future studies and therapeutic avenues. The identification of an immunosuppressive state in high-neural glioblastoma is concordant with recent findings by Nejo et al. who described immunosuppressive mechanisms in thrombospondin-1-upregulated glioma samples^49^. Taken together, stratification of IDH-wildtype gliomas based on their epigenetic neural signature may provide a potential tool for predicting response to neuroscience-guided therapies.

## Conclusion

Overall, the definition of a high-neural signature in IDH-wildtype glioma revealed an OPC- and NPC-like character with a malignant stem cell-like state that affects patient survival, remains stable during therapy, and is conserved in preclinical models. This knowledge supports clinicians in stratifying glioma patients according to their prognosis and determining the surgical and neuro-oncological benefit for current standard of care. Lastly, the here presented clinical translation in the field of glioma neuroscience using an epigenetic neural signature may advance the development of trials with neuroscience-guided therapies.

## METHODS

### DNA Methylation Profiling

DNA was extracted from tumors, extracellular vesicles, and bulk plasma, and analyzed for genome-wide DNA methylation patterns using the Illumina EPIC (850k) array. Processing of DNA methylation data was performed with custom approaches.^50^ Methylation profiling results from first surgery were submitted to the molecular neuropathology (MNP) methylation classifier v12.5 hosted by the German Cancer Research Center (DKFZ).^17^ Patients were included if the calibrated score for the specific methylation class was >0.84 at time of diagnosis in accordance with recommendations by Capper et al.^50^ For *IDH*-wildtype glioblastoma, patients with a score below 0.84 but above 0.7 with a combined gain of chromosome 7 and loss of chromosome 10 or amplification of epidermal growth factor receptor (*EGFR*) were included in accordance with cIMPACT-NOW criteria.^51^ Furthermore, a class member score of ≥ 0.5 for one of the glioblastoma subclasses was required. Evaluation of the *MGMT* promoter methylation status was made from the classifier output v12.5 using the *MGMT*-STP27 method.^52^

### Processing of Methylation Arrays

All idats corresponding to methylation array data were processed similarly using the minfi package in R (version 1.40.0).^53^ The data was processed using the preprocessIllumina function. Only probes with detection p-values <0.01 were kept for further analysis. Also, probes with <3 beads in at least 5% of samples, as well as all non-CpG probes, SNP-related probes, and probes located on X and Y chromosomes were discarded. The CpG intensities were converted into beta values representing total methylation levels (between 0 and 1).

### Cell Type Deconvolution

Non-negative least square (NNLS) linear regression was used in deconvolving the beta values of methylation arrays into cell type components.^16, 54, 55^ As a reference, a publicly available signature was obtained from Moss et al. (2018) consisting of gene expressions for 25 cell type components (Monocytes_EPIC, B-cells_EPIC, CD4T-cells_EPIC, NK-cells_EPIC, CD8T-cells_EPIC, Neutrophils_EPIC, Erythrocyte_progenitors, Adipocytes, Cortical_neurons, Hepatocytes, Lung_cells, Pancreatic_beta_cells, Pancreatic_acinar_cells, Pancreatic_duct_cells, Vascular_endothelial_cells, Colon_epithelial_cells, Left_atrium, Bladder, Breast, Head_and_neck_larynx, Kidney, Prostate, Thyroid, Upper_GI, Uterus_cervix) and 6,105 unique CpGs.^16^

### Integrative Analysis of Methylation and Gene Expression

The analysis of gene expression co-correlation networks was conducted using Weighted Correlation Network Analysis (WGCNA)^56^, in which the epigenetic Moss-signature was incorporated as trait features. Initially, we calculated the optimal soft power to achieve a scale-free topology. This was done by fitting a model for different soft power thresholds (ranging from 1 to 20), with an optimal power of 16. Following this, a signed co-expression network was created utilizing the Topological Overlap Matrix (TOM)^57^ via the hdWGCNA’s ConstructNetwork function^58^. For dimension reduction and visualization of the co-expression network, we employed the Uniform Manifold Approximation and Projection (UMAP) via the ModuleUMAPPlot function. We then identified the hub genes within each module by calculating module connectivity using the ModuleConnectivity function. Gene ontology analysis was subsequently performed on the top 100 module-associated genes using the compareCluster function. Visualization of module-associated pathway activations was accomplished using the clusterProfiler package^59^, specifically via the dotplot function.

### Single Cell Data Analysis

To contextualize the gene expression modules significantly associated with the low-/ high-neural epigenetic phenotype, we computed eigengene signatures and examined their expression patterns using the GBMap single-cell reference dataset^60^. We downloaded and processed GBMap using the Seurat package. The AddModuleScore function of the Seurat package was used to compute the module eigengene score for each cell. For visualization, we projected the model expression onto the cell-level UMAP (Uniform Manifold Approximation and Projection) provided by GBMap’s integration algorithm.

### Spatially Resolved Transcriptomics Data Analysis

We accessed spatial transcriptomic data from our institute recently published^20^ and preprocessed the corresponding EPIC methylation data by the pipeline above. Computational analysis was employed by the SPATA2 package (v2.01). For spatial projection of the module eigengene signatures, we used the joinWithGenes function and averaged the expression across all genes. Spatial surface plots were performed by the plotSurface function without smoothing. Spatial correlation analysis was performed by the MERINGUE package^61^ using the spatial cross correlation analysis. Spatial proximity analysis, we performed a correlation-based analysis using low- or high-neural glioblastoma samples. A spatial correlation matrix was generated using SPATA2’s joinWithFeatures function, which incorporates the annotation_level_4 data from the GBMap single cell deconvolution. The correlation matrices were then averaged using the Reduce function. To estimate the average cellular abundance of each cell type/state, we employed a similar approach. The resulting correlation matrix was transformed into a distance matrix, with correlation values subtracted from 1. We then applied a threshold, setting distances derived from correlations less than 0.5 to zero, effectively removing low correlation connections. Subsequently, we created a graph object from the distance matrix using the graph_from_adjacency_matrix function from the igraph package^62^. We added attribute data (cell type and abundance) to the graph vertices.

Next, we computed a minimum spanning tree from the graph to simplify and highlight the core structure of the network. Edge weights were normalized to a range between 0.5 and 2, setting the basis for edge width in subsequent graph visualization. Finally, we visualized the graph using the ggraph package^63^, incorporating edge links and node points, which were color-coded and sized according to cellular abundance. Node labels were added with the geom_node_text function and repelled for better visibility.

### Cell State Composition Analysis

To infer the abundance of cell type and cell state in the samples, we subjected each sample to bulk DNA methylation assay using EPIC arrays and applied the Silverbush et al. deconvolution method^64^. The deconvolution method is a reference free method that uses a hierarchical matrix factorization approach inferring both cell types and the cell states therein. The method was trained on the DKFZ glioblastoma cohort and tested on TCGA glioblastoma cohort and was able to infer the abundance of cell types in the microenvironment (immune, glia and neuron) and malignant cell states (malignant stem-like cells component and two differentiated cells components). We applied the method as described in Silverbush et al. using the cell type and cell state encoding provided in the manuscript and via the engine provided in EpiDISH^65^ package, with RPC method and maximum iterations of 2000.

### DNA Tumor Purity

Tumor-purity was calculated using the RF_purify Package in R.^66^ This package uses the “absolute” method which measures the frequency of somatic mutations within the tumor sample and relates this to the entire DNA quantity.^67^

### Isolation and Analysis of Extracellular Vesicles (EVs)

EVs were isolated from plasma of glioblastoma patients by differential centrifugation as previously described.^31, 68^ Plasma and culture supernatants were centrifuged at 300 x g for 5 min to eliminate cells, followed by 2000 x g for 10 min to remove platelets and remaining cell debris. Thereafter, the cleared plasma and supernatants were centrifuged at 10,000 x g for 30 min (4°C) to remove large vesicles, and then followed to ultracentrifugation at 100,000 x g for 70 min (4°C), where EV pellets were resuspended with 0.22µm-filtered (Millipore) PBS. The concentration and size of EVs were determined by nanoparticle tracking analysis (NTA), using an LM14 instrument (NanoSight, Malvern Panalytical) equipped with a 638 nm laser and a Merlin F-033B IRF camera (Adept Electronic Solutions). EV-enriched samples were diluted 1:300 in PBS prior to NTA. Triple movies (30 seconds each) were recorded on camera level 15, and then analyzed with detection threshold 6 in NTA 3.2 Build 16. As routine, EVs were also characterized according to size and morphology by electron microscopy, and according to EV markers (CD9, CD63, CD81) by Imaging Flow Cytometry (data not shown). DNA was extracted from EVs using the MasterPure Complete DNA and RNA Purification Kit (Biosearch Technologies). For comparison purposes, bulk cfDNA was isolated from plasma with the MagMax™ cfDNA Isolation Kit (Applied Biosystems).

### Detection of BDNF Serum Levels

Plasma from glioblastoma patients was isolated by double spin centrifugation of whole blood. Samples were aliquoted and stored at -80 C before use. BDNF plasma levels were detected using the LEGENDplex Neuroinflammation Panel 1 (Biolegend, San Diego, CA, USA). Data was acquired using the BD LSR Fortessa and Beckman Coulter Cytoflex LX flow cytometer and analyzed with the BioLegend LEGENDplex software.

### Proteomic Processing of Human Glioblastoma Samples

FFPE samples of tumors were obtained from tissue archives from the neuropathology unit in Hamburg. Tumor samples were fixed in 4 % paraformaldehyde, dehydrated, embedded in paraffin, and sectioned at 10 μm for microdissection using standard laboratory protocols. For paraffin removal FFPE tissue sections were incubated in 0.5 mL n-heptane at room temperature for 30 min, using a ThermoMixer (ThermoMixer^®^ 5436, Eppendorf). Samples were centrifuged at 14.000 g for 5 min and the supernatant was discarded. Samples were reconditioned with 70% ethanol and centrifuged at 14.000 g for 5 min. The supernatant was discarded. The procedure was repeated twice. Pellets were dissolved in 150 µL 1 % w/v sodium deoxycholate (SDC) in 0.1 M triethylammonium bicarbonate buffer (TEAB) and incubated for 1 h at 95 °C for reverse formalin fixation. Samples were sonicated for 5 seconds at an energy of 25% to destroy interfering DNA. A bicinchoninic acid (BCA) assay was performed (Pierce™ BCA Protein Assay Kit, Thermo Scientific) to determine the protein concentration, following the manufacturer’s instructions. Tryptic digestion was performed for 20 ug protein, using the Single-pot, solid-phase-enhanced sample preparation (SP3) protocol^69^. Eluted Peptides were dried in a Savant SpeedVac Vacumconcentrator (Thermo Fisher Scientific, Waltham, USA) and stored at -20° until further use. Directly prior to measurement dried peptides were resolved in 0.1% FA to a final concentration of 1 μg/μl. In total 1 μg was subjected to mass spectrometric analysis.

### Liquid Chromatography–Tandem Mass Spectrometer Parameters

Liquid chromatography–tandem mass spectrometer (LC–MS/MS) measurements were performed on a quadrupole-ion-trap-orbitrap mass spectrometer (MS, QExactive, Thermo Fisher Scientific, Waltham, MA, USA) coupled to a nano-UPLC (Dionex Ultimate 3000 UPLC system, Thermo Fisher Scientific, Waltham, MA, USA). Tryptic peptides were injected to the LC system via an autosampler, purified and desalted by using a reversed phase trapping column (Acclaim PepMap 100 C18 trap; 100 μm × 2 cm, 100 A pore size, 5 μm particle size; Thermo Fisher Scientific, Waltham, MA, USA), and thereafter separated with a reversed phase column (Acclaim PepMap 100 C18; 75 μm × 25 cm, 100 A pore size, 2 μm particle size, Thermo Fisher Scientific, Waltham, MA, USA). Trapping was performed for 5 min at a flow rate of 5 µL/min with 98% solvent A (0.1% FA) and 2% solvent B (0.1% FA in ACN). Separation and elution of peptides were achieved by a linear gradient from 2 to 30% solvent B in 65 min at a flow rate of 0.3 µL/min. Eluting peptides were ionized using a nano-electrospray ionization source (nano-ESI) with a spray voltage of 1800 V, transferred into the MS, and analyzed in data dependent acquisition (DDA) mode. For each MS1 scan, ions were accumulated for a maximum of 240 ms or until a charge density of 1 × 1^6 ions (AGC target) were reached. Fourier-transformation-based mass analysis of the data from the orbitrap mass analyzer was performed by covering a mass range of 400–1200 *m*/*z* with a resolution of 70,000 at *m*/*z* = 200. Peptides with charge states between 2+–5+ above an intensity threshold of 5 000 were isolated within a 2.0*m*/*z* isolation window in top-speed mode for 3 s from each precursor scan and fragmented with a normalized collision energy of 25%, using higher energy collisional dissociation (HCD). MS2 scanning was performed, using an orbitrap mass analyzer, with a starting mass of 100 m/z at an orbitrap resolution of 17,500 at *m*/*z* = 200 and accumulated for 50 ms or to an AGC target of 1 × 105^. Already fragmented peptides were excluded for 20 s.

### Proteomic Data Processing

Proteomic samples (n=28) were measured with liquid chromatography tandem mass spectrometry (LC-MS/MS) systems and processed with Proteome Discoverer 3.0. and searched against a reviewed FASTA database (UniProtKB: Swiss-Prot, Homo sapiens, February 2022, 20300 entries). To cope with protein injection amount differences, the protein abundances were normalized at the peptide level. Perseus 2.0.3 was used to obtain log2 transformed intensities. The imputation was performed using the Random Forest imputation algorithm (Hyperparameters: 1000 Trees and 10 repetitions) in RStudio 4.3.

### Weighted Correlation Network Analysis (WGCNA)

The WGCNA package in R (version 1.70.3) was used to identify gene co-expression gene modules.^56^ The minimum module size was set to 10 and a merging threshold of 0.40 was defined. Based on the assessment of scale-free topology, soft-power of 9 was selected. To construct modules, we first corrected for any technical batch effect using Empirical Bayes-moderated adjustment using empiricalBayesLM function of WGCNA. Modules were assessed based on their correlation with traits (low and high) and their levels of significance (associated with two-tailed Student’s t-test). The significant modules (p<0.05) were used for further analysis. All genesets within a module were used for overrepresentation analysis using clusterProfiler package^59^ in R (Version 4.2.0). Further to identify cell type enrichment within each module, gene-sets from PanglaoDB3 were used through enrichr in python (Package maayanlab_bioinformatics, version 0.5.4)^70^. To assess the module scores on single-cells, Scanpy’s score_genes function was used to calculate module scores using core glioblastoma single-cell atlas^60^.

### Mice Housing

*In vivo* experiments were conducted in accordance with protocols approved by the Stanford University Institutional Animal Care and Use Committee (IACUC) as well as the University Medical Center Hamburg-Eppendorf (Hamburg, Germany). Experiments were performed in accordance with institutional guidelines and explicit permission from the local authorities (Behörde für Soziales, Gesundheit und Verbraucherschutz Hamburg, Germany). Animals were housed according to standard guidelines under pathogen-free conditions, in temperature- and humidity-controlled housing with free access to food and water in a 12 h light:12 h dark cycle. For brain tumor xenograft experiments, the IACUC does not set a limit on maximal tumor volume but rather on indications of morbidity. In no experiments were these limits exceeded as mice were euthanized if they exhibited signs of neurological morbidity or if they lost 15% or more of their body weight.

### Orthotopic Xenografting of Patient-Derived Low- and High-Neural Glioblastoma Cells

For xenograft studies as presented in Fig. 4g-m, NSG mice (NOD-SCID-IL2R gamma chain-deficient, The Jackson Laboratory) were used, and experiments were performed at the Stanford University (United States). Male and female mice were used equally. A single-cell suspension from cultured primary patient-derived low- (“UCSF-UKE-1”) or high-neural (“UCSF-UKE-2”) glioblastoma neurospheres was prepared in sterile HBSS immediately before the xenograft procedure. Mice at postnatal day (P) 28–30 were anaesthetized with 1– 4% isoflurane and placed in a stereotactic apparatus. The cranium was exposed through midline incision under aseptic conditions. Approximately 150,000 cells in 3 μl sterile HBSS were stereotactically implanted into the premotor cortex (M2) through a 26-gauge burr hole, using a digital pump at infusion rate of 1.0 μl min^−1^. Stereotactic coordinates used were as follows: 0.5 mm lateral to midline, 1.0 mm anterior to bregma, −1.0 mm deep to cortical surface.

Mice survival data from the orthotopic xenografts demonstrated in Fig. 4e were performed on NMRI-Foxn1nu immunodeficient mice (Janvier-Labs) and conducted at the University Medical Center Hamburg-Eppendorf (Germany). After dissociation, neurospheres from cultured primary patient-derived low- (“GS-8”, “GS-10”, “GS-73”, and “GS-80”) or high-neural (“GS-57”, “GS-74”, “GS-75”, “GS-101”) glioblastoma were resuspended in a concentration of 100.000 cells/ul in HBSS and 2ul was injected in the striatum at the following stereotactic coordinates as follows: 2.0mm lateral to Bregma, 1.0mm anterior to Bregma, and -2.8mm deep to cortical surface. Cells were implanted using a Hamilton syringe with a 30-gauge needle. Further data is available in extended data 5.

### Perfusion and Immunofluorescence Staining

Eight weeks after xenograft, low and high neural glioblastoma-bearing mice were anaesthetized with intraperitoneal avertin (tribromoethanol), then transcardially perfused with 20 ml of PBS. Brains were fixed in 4% PFA overnight at 4 °C, then transferred to 30% sucrose for cryoprotection for 48 h. Brains were then embedded in Tissue-Tek O.C.T. (Sakura) and sectioned in the coronal plane at 40 μm using a sliding microtome (Microm HM450; Thermo Scientific). For immunofluorescence, coronal sections were incubated in blocking solution (3% normal donkey serum, 0.3% Triton X-100 in TBS) at room temperature for 30 min. Mouse anti-human nuclei clone 235-1 (1:100; Millipore), rabbit anti-Ki67 (1:500; Abcam ab15580), rat anti-MBP (1:200; Abcam ab7349), mouse anti-nestin (1:500; Abcam ab6320), guinea pig anti-synapsin1/2 (1:500; Synaptic Systems), chicken anti-neurofilament (M+H; 1:1000; Aves Labs) or PSD95 (1:500, Abcam ab18258), were diluted in antibody diluent solution (1% normal donkey serum in 0.3% Triton X-100 in TBS) and incubated overnight at 4 °C. Sections were then rinsed three times in TBS and incubated in secondary antibody solution (Alexa 488 donkey anti-rabbit IgG; Alexa 594 donkey anti-mouse IgG, Alexa 647 donkey anti-chicken IgG, Alexa 405 donkey anti-guinea pig IgG, Alexa 647 donkey anti-rabbit IgG, or Alexa 594 donkey anti-mouse IgG all used at 1:500 (Jackson Immuno Research) in antibody diluent at 4 °C. Sections were rinsed three times in TBS and mounted with ProLong Gold Mounting medium (Life Technologies).

### Confocal Imaging and Quantification of Cell Proliferation and Tumor Burden

Cell quantification within xenografts was performed by a blinded investigator using live counting on a 20x objective of a Zeiss LSM900 scanning confocal microscope and Zen 3.7 imaging software (Carl Zeiss). For overall tumor burden analysis, a 1-in-6 series of coronal brain sections were selected with 4 consecutive slices (4 fields per slice) at approximately 1.1–0.86 mm anterior to bregma analysed. Within each field, all HNA-positive tumor cells were quantified to determine tumor burden within the areas quantified. HNA-positive tumor cells were then assessed for co-labelling with Ki67. To calculate the proliferation index (the percentage of proliferating tumor cells for each mouse), the total number of HNA-positive cells co-labelled with Ki67 across all areas quantified was divided by the total number of cells counted across all areas quantified (Ki67^+^/ HNA^+^).

### Confocal Puncta Quantification

Images were collected using a 63×oil-immersion objective on a Zeiss LSM900 confocal microscope. Colocalization of all synaptic puncta images from low and high-neural glioblastoma xenograft samples described above were analyzed using a custom ImageJ processing script written at the Stanford Shriram Cell Science Imaging Facility to define each pre- and postsynaptic puncta and determine colocalization within a defined proximity of 1.5 μM. To partially subtract local background, we used the ImageJ rolling ball background subtraction (https://imagej.net/Rolling_Ball_Background_Subtraction). The peaks were found using the imglib2 DogDetection plugin (https://github.com/imglib/imglib2algorithm/blob/master/src/main/java/ net/imglib2/ algorithm/dog/DogDetection.java). In this plugin, the difference of Gaussians is used to enhance the signal of interest using two different sigmas: a ‘smaller’ sigma, which defines the smallest object to be found and a ‘larger’ sigma, for the largest object. The plugin then identifies the objects that are above the min peak value and assigns regions of interest (ROIs) to each channel. The number of neuron and glioma ROIs are counted, and the script extracts the number of glioma ROIs within 1.5μm of the neuron ROIs. This script was implemented in Fiji/ImageJ using the ImgLib2 and ImageJ Ops (https://imagej.net/ImageJ_Ops) libraries.

### Sample Preparation and Image Acquisition for Electron Microscopy

Twelve weeks after xenografting of low- (n =3, “UCSF-UKE-1”) and high-neural glioblastoma cell (n = 3, “UCSF-UKE-2”), mice were euthanized by transcardial perfusion with Karnovsky’s fixative: 2% glutaraldehyde (EMS, 16000) and 4% PFA (EMS, 15700) in 0.1 M sodium cacodylate (EMS, 12300), pH 7.4. Transmission electron microscopy (TEM) was performed in the tumor mass within the CA1 region of the hippocampus for all xenograft analysis. The samples were then post-fixed in 1% osmium tetroxide (EMS, 19100) for 1 h at 4 °C, washed three times with ultrafiltered water, then en bloc stained overnight at 4 °C. The samples were dehydrated in graded ethanol (50%, 75% and 95%) for 15 min each at 4 °C; the samples were then allowed to equilibrate to room temperature and were rinsed in 100% ethanol twice, followed by acetonitrile for 15 min. The samples were infiltrated with EMbed-812 resin (EMS, 14120) mixed 1:1 with acetonitrile for 2 h followed by 2:1 EMbed-812:acetonitrile overnight. The samples were then placed into EMbed-812 for 2 h, then placed into TAAB capsules filled with fresh resin, which were then placed into a 65 °C oven overnight. Sections were taken between 40 nm and 60 nm on a Leica Ultracut S (Leica) and mounted on 100-mesh Ni grids (EMS FCF100-Ni). For immunohistochemistry, microetching was done with 10% periodic acid and eluting of osmium with 10% sodium metaperiodate for 15 min at room temperature on parafilm. Grids were rinsed with water three times, followed by 0.5 M glycine quench, and then incubated in blocking solution (0.5% BSA, 0.5% ovalbumin in PBST) at room temperature for 20 min. Primary goat anti-RFP (1: 300, ABIN6254205) was diluted in the same blocking solution and incubated overnight at 4 °C. The next day, grids were rinsed in PBS three times, and incubated in secondary antibodies (1:10 10 nm gold-conjugated IgG, TED Pella, 15796) for 1 h at room temperature and rinsed with PBST followed by water. For each staining set, samples that did not contain any RFP-expressing cells were stained simultaneously to control for any non-specific binding. Grids were contrast stained for 30 s in 3.5% uranyl acetate in 50% acetone followed by staining in 0.2% lead citrate for 90 s. The samples were imaged using a JEOL JEM-1400 TEM at 120 kV and images were collected using a Gatan Orius digital camera.

### Electron Microscopy Data Analysis

Sections from xenografted hippocampi of mice were imaged using TEM imaging. The xenografts were originally generated for a study by Krishna et al.^10^ and mouse tissue was re-analyzed after epigenetic profiling and assignment to low- or high-neural glioblastoma groups. Here, 42 sections of high-neural glioblastoma across 3 mice and 45 sections of low-neural glioblastoma across 3 mice were analyzed. Electron microscopy images were taken at 6,000× with a field of view of 15.75 μm^2^. Glioma cells were counted and analyzed after identification of immunogold particle labelling with three or more particles. Furthermore, to determine synaptic structures all three of the following criteria had to be clearly met as previously described^3^: 1) presence of synaptic vesicle clusters, 2) visually apparent synaptic cleft, and 3) identification of postsynaptic density in the glioma cell. To quantify the percentage of glioma cells forming synaptic structures, the number of glioma-to-neuron synapses identified was divided by the total number of glioma cells analyzed.

### Cell Culture

Fresh glioblastoma samples were obtained from patients operated in the Department of Neurosurgery, University Medical Center Hamburg-Eppendorf (Germany). Samples were immediately placed in Hankś balanced salt solution (HBSS, Invitrogen), transferred to the laboratory and processed within 20 min. The tissue was cut into <1 mm^3^ fragments, washed with HBSS and digested with 1 mg/ml collagenase/dispase (Roche) for 30 min at 37 °C. Digested fragments were filtered using a 70 μm cell mesh (Sigma-Aldrich), and the cells were seeded into T25 flasks at 2500–5000 cells/cm^2^. The culture medium consisted of neurobasal medium (Invitrogen) with B27 supplement (20 μl/ ml, Invitrogen), Glutamax (10 μl/ ml, Invitrogen), fibroblast growth factor-2 (20 ng/ml, Peprotech), epidermal growth factor (20 ng/ml, Peprotech) and heparin (32 IE/ml, Ratiopharm). Growth factors and heparin were renewed twice weekly. Spheres were split by mechanical dissociation when they reached a size of 200–500 μm. In this study analyzed cell cultures with clinical data are represented in extended data 4. Long-term cultivation cell cultures were used from a publically available data set (n = 7, GSE181314) and one in house cell line (n = 1).

### Neuron-Glioma Co-Culture Experiments

Neurons were isolated from CD1 (The Jackson Laboratory) mice at P0 using the Neural Tissue Dissociation Kit - Postnatal Neurons (Miltenyi), and followed by the Neuron Isolation Kit, Mouse (Miltenyi). After isolation, 150.000 neurons were plated onto glass coverslips (Electron Microscopy Services) after pre-treatment with poly-l-lysine (Sigma) and mouse laminin (Thermo Fisher) as described previously^3^. Neurons are cultured in BrainPhys neuronal medium (StemCell Technologies) containing B27 (Invitrogen), BDNF (10ng ml^-1^, Shenandoah), GDNF (5ng ml^-1^, Shenandoah), TRO19622 (5μM; Tocris), β-mercaptoethanol (Gibco). Half of the medium was replenished on days *in vitro* (DIV) 1 and 3. On DIV 5, half of the medium was replaced in the morning. In the afternoon, the medium was again replaced with half serum-free medium containing 75.000 cells from patient-derived low- (“UCSF-UKE-1”) or high-neural (“UCSF-UKE-2”) cell cultures. Cells were cultured with neurons for 72 h and then fixed with 4% paraformaldehyde (PFA) for 20 min at room temperature and stained for puncta quantification as described above.

### EdU Proliferation Assay

For EdU proliferation assays, coverslips were prepared as described above. Again, at DIV 5, low-neural (“UCSF-UKE-1”) or high-neural (“UCSF-UKE-2”) glioblastoma cells were added to the neuron cultures. Forty-eight hours after addition of glioblastoma cells, slides were treated with 10 μM EdU. Cells were fixed after an additional 24 h using 4% PFA and stained using the Click-iT EdU kit and protocol (Invitrogen). Proliferation index was then determined by quantifying the percentage of EdU labelled glioblastoma cells (identified by EdU^+^/DAPI^+^) over total number of glioblastoma cells using confocal microscopy.

### 3D Migration Assay

3D migration experiments were performed as previously described (Vinci et al., Methods Mol. Biol. 2013) with some modifications. Briefly, 96-well flat-bottomed plates (Falcon) were coated with 2.5μg per 50μl laminin per well (Thermo Fisher) in sterile water. After coating, a total of 200μl of culture medium per well was added to each well. A total of 100μl of medium was taken from 96-well round bottom ULA plates containing ∼200μm diameter neurospheres of low- (“UCSF-UKE-1”) and high-neural (“UCSF-UKE-2”) glioblastoma lines, and the remaining medium including neurospheres was transferred into the pre-coated plates. Images were then acquired using an Evos M5000 microscope (Thermo Fisher Scientific) at time zero, 24, 48, and 72 hours after encapsulation. Image analysis was performed using ImageJ by measuring the diameter of the invasive area. The extent of cell migration on the laminin was measured for six replicate wells normalized to the diameter of each spheroid at time zero and the data is presented as a mean ratio for three biological replicates.

### Patient Cohorts

In this study, several patients’ cohorts depending on the glioma subclass were analyzed. First, a clinical cohort of 363 patients who underwent *IDH*-wildtype glioblastoma resection at University Medical Center Hamburg-Eppendorf, University Hospital Frankfurt, or Charité University Hospital Berlin (all Germany) was analyzed. Informed written consent was obtained from all patients and experiments were approved by the medical ethics committee of the Hamburg chamber of physicians (PV4904). Second, we included patients from the GBM-TCGA cohort for external validation^18^. Third, a clinical cohort of pediatric and adolescent patients who underwent surgery for *H3 K27*-altered DMG at University Medical Center Hamburg-Eppendorf (Germany) was established and extended with two cohorts from previously published studies by Sturm et al. and Chen et al.^29, 36^. Last, the reference and diagnostic set (n=3905) published by Capper et al. was used for deconvolution analyses^17^.

### Clinical Definitions

For the internal clinical patients cohort, diagnosis was based on the WHO classification.^71^ The extent of resection (EOR) was stratified into gross total resection (GTR), near GTR, and partial resection (PR). A GTR was defined as a complete removal of contrast-enhancing parts, a near GTR as a removal of more than 90% of the contrast-enhancing parts, whereas a resection of lower than 90% was defined as PR/biopsy. The EOR of contrast-enhancing parts was evaluated by MRI performed up to 48 h after index surgery. Overall survival (OS) was calculated from diagnosis until death or last follow-up, and progression-free survival (PFS) from diagnosis until progression according to Response Assessment in Neuro-Oncology (RANO) criteria based on local assessment^72^. Seizures and use of antiepileptic medication were defined according to the current guidelines of the International League Against Epilepsy (ILAE)^73^. For 3D volumetric segmentation, we analyzed T1-weighted as well as T2-weighted FLAIR (fluid attenuated inversion recovery) magnetic resonance imaging (MRI) axial images of glioblastoma patients before surgery. The program BRAINLAB was used for all analyses. To measure tumor volume, the tumor region of interest was delineated with the tool “Smart Brush” in every slice by hand, enabling a multiplanar 3D reconstruction. With this methodology, the volume of contrast enhancement, FLAIR hyperintensity, and necrotic volume was assessed in cm^3^.

### Stereotactic Biopsies for Spatial Sample Collection

Biopsies were obtained using a cranial navigation system (Brainlab AG, Munich, Germany) and intraoperative neuronavigation. To limit the influence of brain shift, biopsies were obtained before tumor removal at the beginning of surgery with minimal dural opening. Tissue samples were then transferred to 10% buffered formalin and sent to the Department of Neuropathology for further processing and histopathological evaluation.

### Measurement of Functional Connectivity using Magnetoencephalography

Tumor tissues with high (HFC) and low (LFC) functional connectivity sampled during surgery based on preoperative magnetoencephalography (MEG) were obtained from *IDH*-wildtype glioblastoma patients operated in the Department of Neurosurgery, University of California, San Francisco as described previously^10^. From each formalin-fixed paraffin-embedded (FFPE) tissue block, 4 serial sections of an approximate thickness of 10 µm (in total 40 µm) were used for DNA extraction. DNA was extracted with the QIAamp DNA FFPE Kit™ (QIAGEN). DNA was quantified using the Nanodrop Spectrophotometer (Thermo Scientific). The ratio of OD at 260 nm to OD 280 nm was calculated and served as criteria for DNA quality.

### Functional Connectivity by Resting-State Functional Magnetic Resonance Imaging

44 treatment-naîve glioblastoma patients (mean age: 65±9 years) underwent resting-state functional magnetic resonance imaging (rsfMRI) before surgery and tumor tissues were analyzed for genome-wide DNA methylation patterns using the Illumina EPIC (850k) array. Functional data were preprocessed using SPM12^74^ as implemented in Matlab 9.5 according to an imaging protocol that was similarly applied and described in previous publications^75, 76^. Briefly, functional images were realigned to the mean functional volume, unwarped and coregistrated to the structural image. Structural and functional images were segmented, bias corrected and spatially normalized (multi-spectral classification), and functional images were smoothed with a 5 mm FWHM Gaussian kernel. Functional images were then slice-time corrected, movement-related time series were regressed out with ICA-AROMA^77^, and data were high-pass filtered (> 0.01 Hz). Contrast-enhancing tumor lesions were segmented semi-automatically using the ITK-SNAP software version 3.4.0^78^ and used as region of interest (ROI) to perform a seed-based correlation analysis and compute the voxel-based tumor to peritumoral connectivity (Fisher z transformation). A 10mm peritumoral distance mask was created by dilating the tumor mask by 10mm and subtracting the tumor area. The mean functional connectivity between tumor and its 10mm peritumoral surrounding was computed using a ROI-to-voxel approach.

### Drug Sensitivity Analysis

The patient-derived low-neural spheroid glioblastoma cell lines GS-11, GS-73, GS-84 and GS-110 as well as the high-neural ones GS-13, GS-74, GS-80, GS-90 and GS-101 (Extended Data 6) were dissociated into single cells and were seeded in Neurobasal medium supplemented with B27, 1% Glutamine, 1% Pen/Strep, 1uL/mL Heparin and 20 ng/mL human FGF and EGF at 1250-7500 cells/well into a clear-bottom, tissue-culture treated 384-well plate (Perkin Elmer, Waltham, Massachusetts, USA). The cells were treated in triplicates with 27 drugs and with DMSO as a control for 48 hours at 37°C and 5% CO2. Afterwards the cells were fixed with 4% PFA (Sigma/Aldrich), blocked with PBS containing 5% FBS, 0.1%TritonX and DAPI (4 ug/mL, #422801, Biolegend) for one hour at room temperature and were stained with vimentin (#677809, Biolegend), cleaved Caspase 3 (#9604S, Cell Signaling) and TUBB3 (#657406, Biolegend) antibodies overnight at 4°C. The plate was imaged with an Opera Phenix automated spinning-disk confocal microscope in three z-stacks at 10x magnification (Perkin Elmer). The maximal intensity projection of the z-stacks was used for segmentation of the spheroids based on their DAPI staining using CellProfiler 2.2.0. Downstream image analysis was performed with MATLAB R2021b. Marker positive cells/spheroids were identified by a linear threshold on the respective channel. The cell counts as well as the average cell/spheroid areas were averaged per condition and compared between drug treatment and the control group.

### Statistical Analysis

Gaussian distribution was confirmed by the Shapiro-Wilk normality test. For parametric data, unpaired two-tailed Student’s t-test or one-way ANOVA with Tukey’s post hoc tests to examine pairwise differences were used as indicated. Survival curves were visualized as results from the Kaplan-Meier method applying two-tailed log rank analyses for analyzing statistical significance. Multivariate analysis for OS and PFS displaying hazard ratios (HRs), and 95% confidence interval (CI) were computed for each group using Cox proportional hazards regression model. All variables associated with OS or PFS with p-value less than 0.05 in univariate analysis were included in the multivariable model. In general, a p-value less than 0.05 was considered statistically significant for all experiments. Statistical analyses and data illustrations were performed using GraphPad Prism 10. Alluvial plots were graphed with R studio.

## Acknowledgement

We thank Lotte Stegat (Department of Neuropathology, University Medical Center Hamburg-Eppendorf, Germany) for contributing data to the DMG cohort. The authors acknowledge Sabine Wutke (University Medical Center Hamburg-Eppendorf, Germany) for graphical assistance. Initial drafts of figures 1a, 4g, and 4m were made with BioRender.com. Lastly, we thank all the patients who gave informed consent and without whom this research would not have been possible.

## Data Availability Statement

Idat files of the clinical cohort (363 glioblastoma patients) will be made available on Gene Expression Omnibus (GEO) prior to publication. The methylation data provided by Capper et al. as illustrated in Supplementary figure 1 are accessible under GSE109381. TCGA-GBM cohort analyzed for external validation and as shown in Figure 1d is accessible under https://portal.gdc.cancer.gov/projects/TCGA-GBM. All other data are available in the article, source data, or from the corresponding author upon reasonable request.

## Authors Contributions

R.D. and F.L.R. designed, conducted, and interpreted all experiments and analyses. R.K. F.H.

T.H. S.B and S.H. performed and analyzed deconvolution, copy number variation and proteomic analysis. M.M. and A.S.R. performed immunoassays quantification of BDNF serum levels. F.L.R., C.M., A.S. and K.L. contributed to cell culture, and extracellular vesicle experiments. A.K.W. and U.S. contributed to DMG cohorts. H.B. calculated DNA tumor purity.

J.N. conducted mass spectrometry proteomic profiling. K.J. and D.D. contributed to functional connectivity measured by resting state MRI. R.D., T.S., L.D., Y.Z., M.W., F.L.R., K.W., P.N.H., D.C., J.O., and P.V. contributed glioblastoma cohorts of each institution. B.W. and J.G. performed stereotactic biopsies for spatial sample collection of human glioblastoma patients.

M.M. contributed single-cell RNA sequencing data of DMG and provided equipment for *in vivo* analyses. R.D. and M.B.K. conducted *in vivo* experiments for analyzing tumor burden and puncta synapse quantification and C.M. performed xenografting for survival analysis. R.D. performed co-culture experiments and migration assays. L.N. performed electron microscopy images which were evaluated by R.D.. D.S., V.H., and M.L.S. performed cell state composition analysis. S.K. and S.H.J. contributed to functional connectivity measured by MEG. D.H.H. performed spatial transcriptomics. M.W., B.S., A.B., and T.W. conducted drug sensitivity analysis. R.D. and F.L.R. wrote the manuscript. All authors contributed to manuscript editing and approved the final manuscript version.

---

**Supplementary figure 1:** Neural signature in different central nervous system tumor entities (left) and healthy brain tissues (right) obtained from the Capper dataset^37^.

**Supplementary figure 2:** Neural signature of all glioblastoma samples. Red line indicates median neural score of all 1058 included glioblastoma patients and defines the cut-off for stratification into low- and high-neural glioblastoma.

**Supplementary figure 3:** Survival analysis of glioblastoma patients applying brain tumor-related cell signatures of the Moss signature.

*OS: overall survival*

**Supplementary figure 4: High-neural glioblastoma is linked with synapse formation and trans-synaptic signaling from methylation and proteomic profiling.**

**a.)** Volcano plot showing differentially methylated CpG sites of genes of the invasivity signature, neuronal signature, and trans-synaptic signaling signature in high-neural glioblastoma.

**b.)** Correlation between neural signature and DNA tumor purity in glioblastoma samples from the clinical cohort.

**c.) – i.) Proteomic profiling of low- and high-neural glioblastoma.**

**c.)** WGCNA analysis showed differentially abundant proteome modules between both neural subgroups.

**d.)** High-neural glioblastomas are clustered to module “blue” (top figure), while low-neural glioblastomas have a higher abundance in module “brown” (bottom figure)

**e.) – f.)** Network analysis revealed **e.)** most expressed proteins and **f.)** associated gene ontology terms for each neural subgroup (high-neural: top, low-neural: bottom).

**g.)** Integrating transcriptomic single-cell data showed an OPC-/NPC-like character in high-neural tumors (“ME blue”).

**h.)** Transcriptomic single-cell CNV plot analysis of glioblastomas with a high-neural signature.

**Supplementary figure 5:** Copy number variation plots for **a.)** all glioblastoma samples and **b.) – c.)** neural subgroups of the clinical cohort (n=363).

**Supplementary figure 6**: Drug sensitivity analysis of low- and high-neural glioblastoma cells.

**a.)** Representative microscopic images for high- (left image) and low-neural (right image) glioblastoma cells. *Green: Vimentin, yellow: cleaved caspase 3, TUBB3: red, DAPI: blue. Scale bars: 10μm*.

**b.)** Drug sensitivity of low- and high-neural glioblastoma cells measured by cleaved caspase 3.

**c.)** Drug sensitivity of low- and high-neural glioblastoma cells measured by average cell area.

**d.)** Statistical difference of sensitivity to various drugs between low- and high-neural glioblastoma cells.

**Supplementary figure 7:** Survival outcome categorized after RANO categories for extent of resection in glioblastoma patients treated by radiochemotherapy with a low- and high-neural signature. *Class 1: 0 cm*^3^ *CE + ≤5 cm*^3^ *nCE tumor, Class 2: ≤1 cm*^3^ *CE, Class 3A: ≤5 cm*^3^ *CE, Class 3B: ≥5 cm*^3^ *CE.*^19^

**Supplementary figure 8: Relevance of neural classification in pediatric and adolescent patients diagnosed with *H3 K27*-altered diffuse midline glioma (DMG).**

**a.)** Association of tumor location with neural signature.

**b.)** Volcano plot showing differentially methylated CpG-sites of genes of the invasivity signature, neuronal signature, and trans-synaptic signaling signature.

**c.)** Cell state composition analysis in low- and high-neural DMG.

**d.)** Synaptic gene expression (*PTPRS*, *ARHGEF2*, *GRIK2*, *DNM3*, *LRRTM2*, *GRIK5*, *NLGN4X*, *NRCAM*, *MAP2*, *INA*, *TMPRSS9*)^6^ is significantly correlated with the stem cell-like state of DMG cells calculated by an overlap of single cell DNA methylation and single cell RNA sequencing (599 cells from 3 study participants) measurements.

**e) – h.)** Kaplan-Meier survival analysis of 72 DMG patients under 18 years of age with a low- and high-neural DMG.

## TABLE LEGENDS

**Supplementary table 1:**

Clinical characteristics of patients with glioblastoma who were treated with combined radio chemotherapy after surgical resection.

*SD: standard deviation, MGMT: O6-methylguanine-DNA-methyltransferase*

**Supplementary table 2:**

Uni- and multivariate analysis of overall survival in patients with glioblastoma.

*HR: hazard ratio, CI: confidence interval, Ref: reference, KPS: Karnofsky Performance Scale, GTR: gross total resection, MGMT: O6-methylguanine-DNA-methyltransferase, CE: contrast-enhancement, FLAIR: fluid attenuated inversion recovery, RTK: receptor tyrosine kinase, MES: mesenchymal*.

**Supplementary table 3:**

Uni- and multivariate analysis of progression-free survival in patients with glioblastoma.

*HR: hazard ratio, CI: confidence interval, Ref: reference, KPS: Karnofsky Performance Scale, GTR: gross total resection, MGMT: O6-methylguanine-DNA-methyltransferase, CE: contrast-enhancement, FLAIR: fluid attenuated inversion recovery, RTK: receptor tyrosine kinase, MES: mesenchymal*.

