## Supplementary Figure for "Epigenetic neural glioblastoma enhances synaptic integration and predicts therapeutic vulnerability"

**Supplementary figure 1**

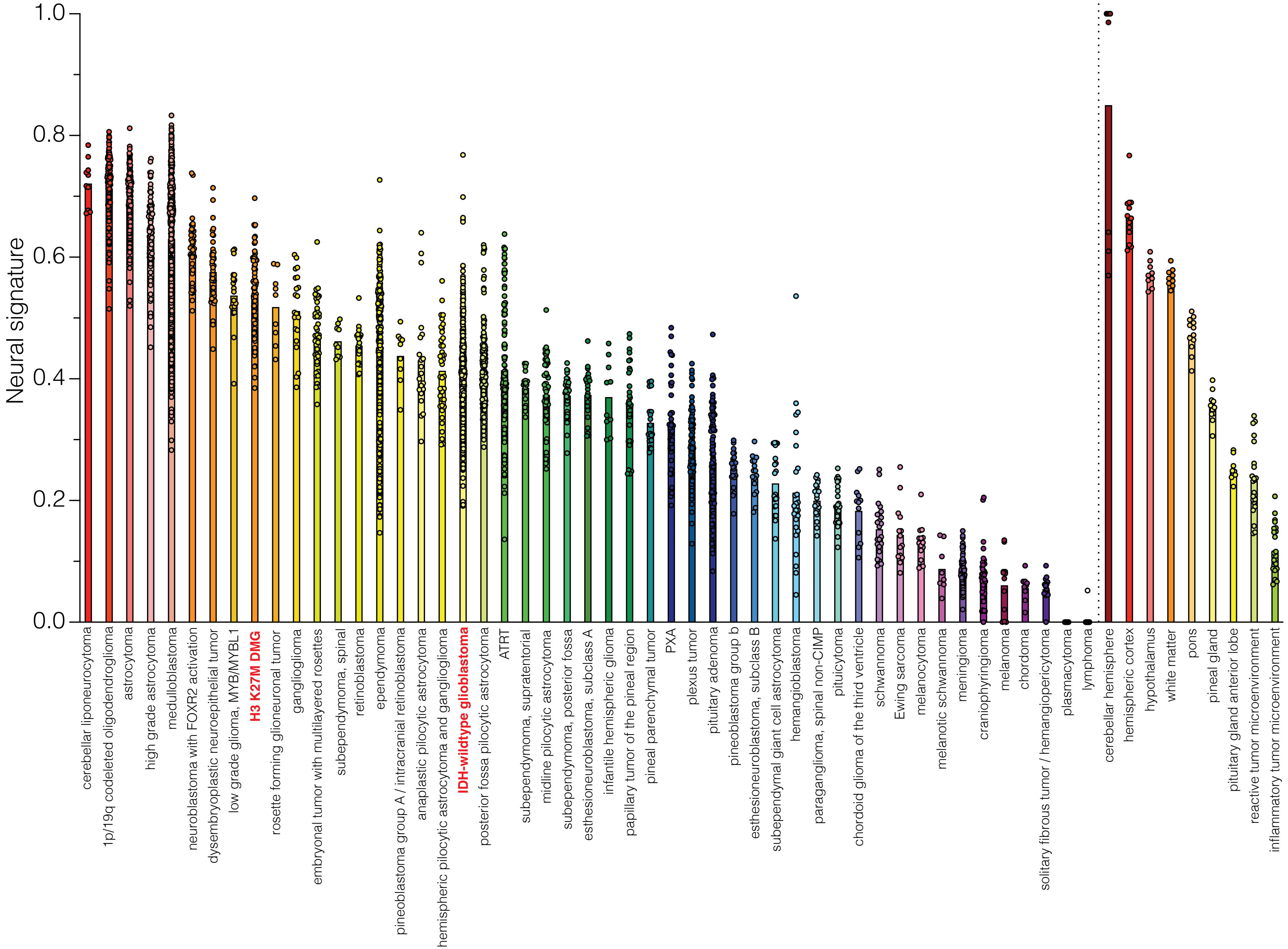

**Supplementary figure 2**

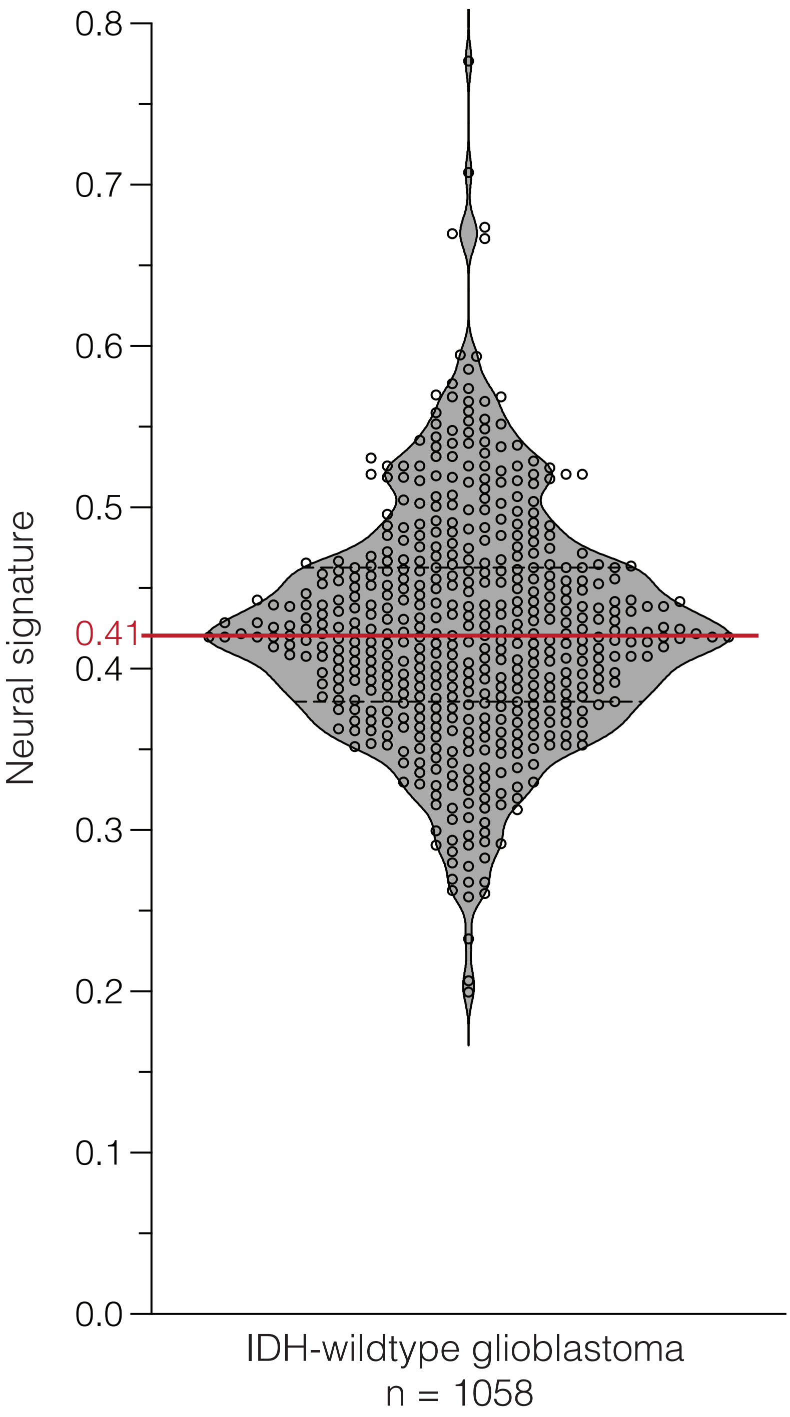

### Supplementary figure 3

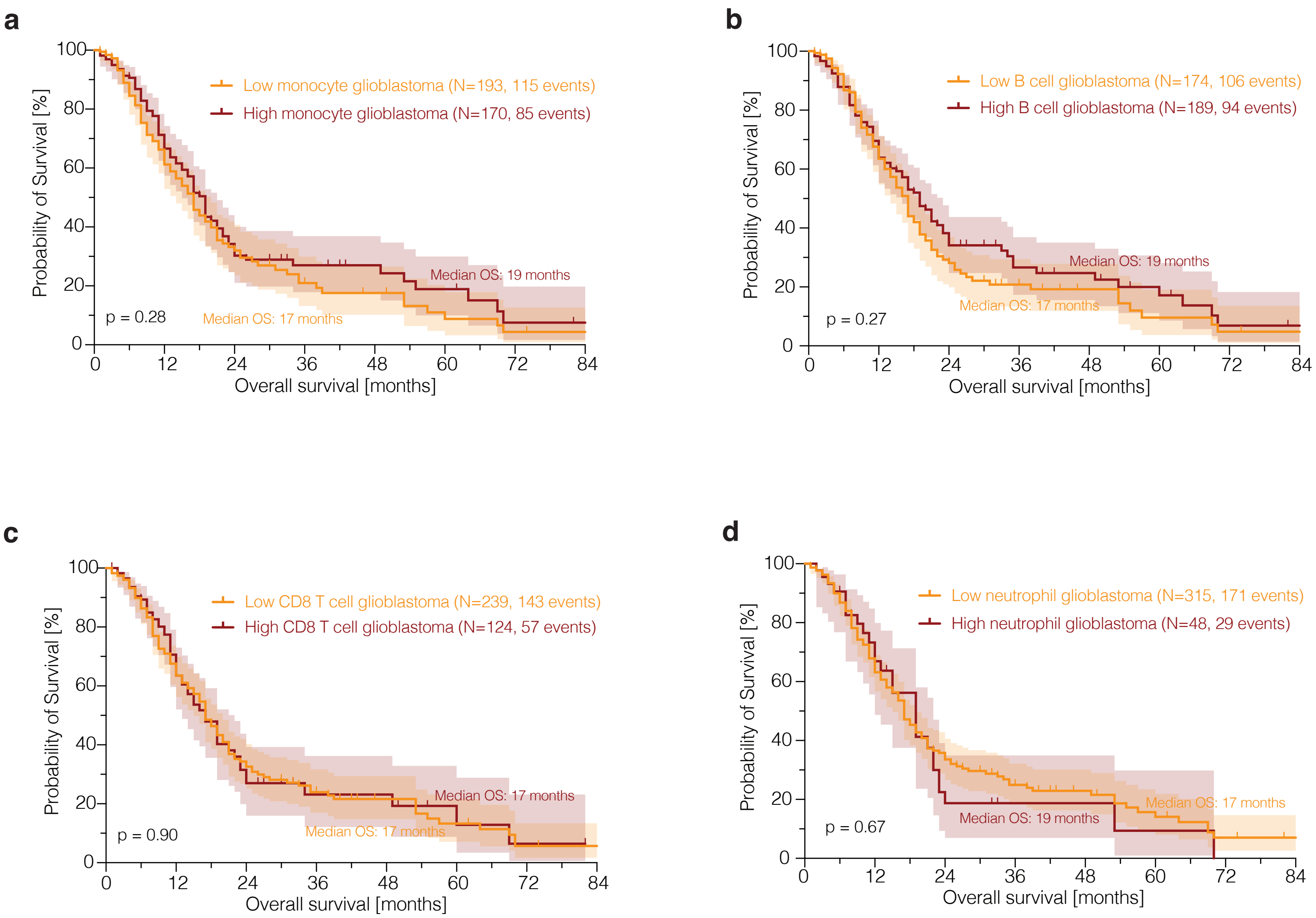

### Supplementary figure 4

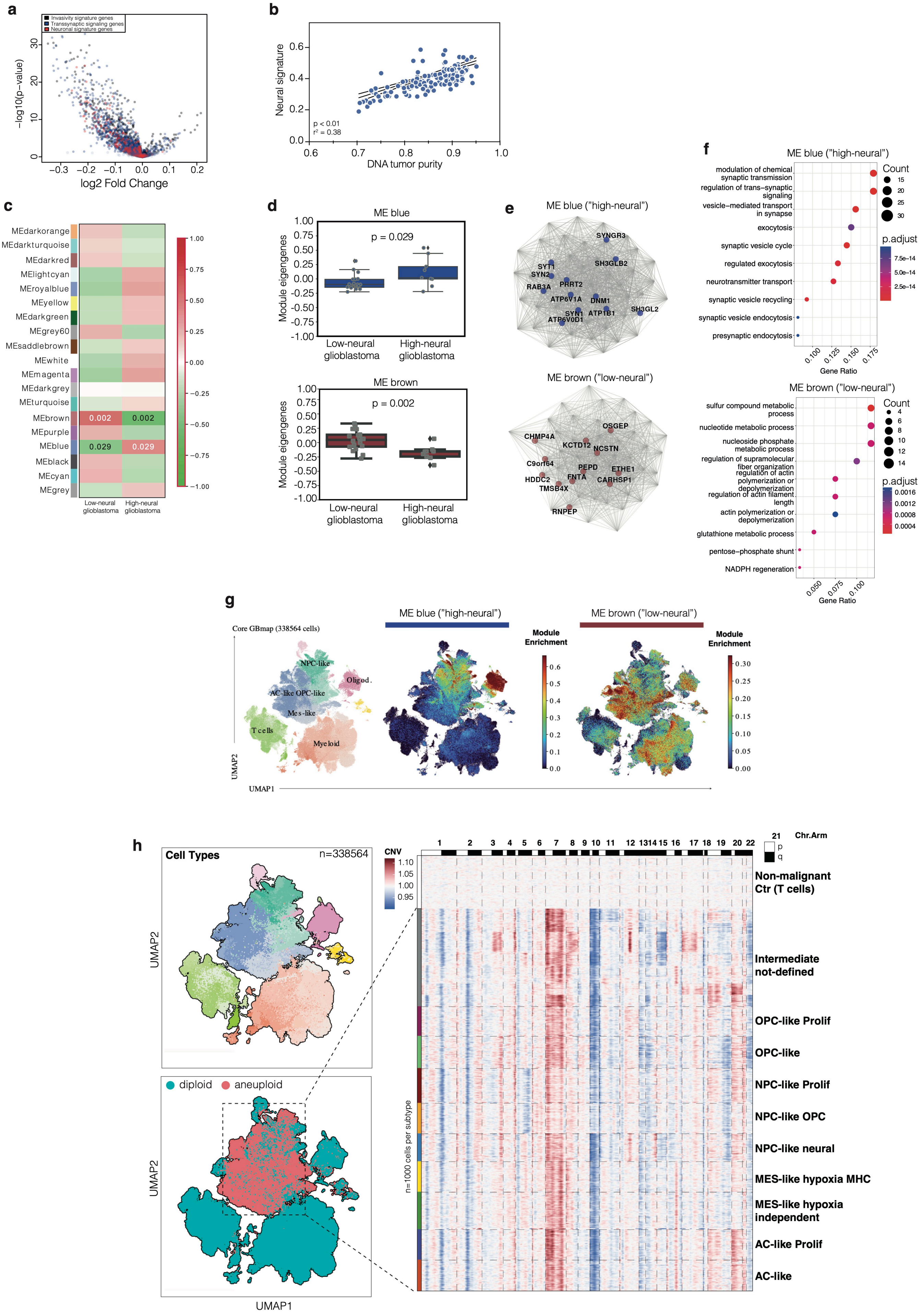

#### Supplementary figure 5

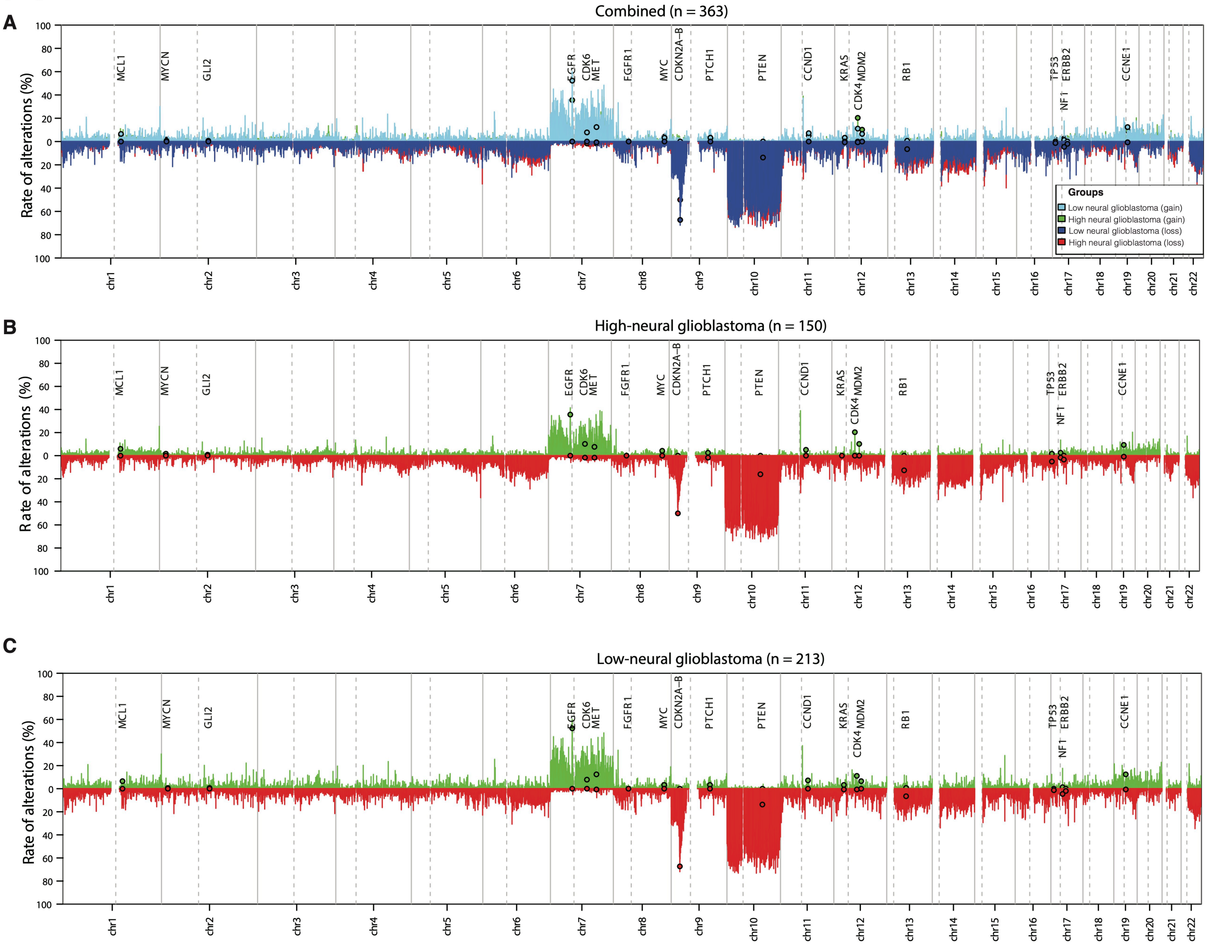

#### Supplementary figure 6

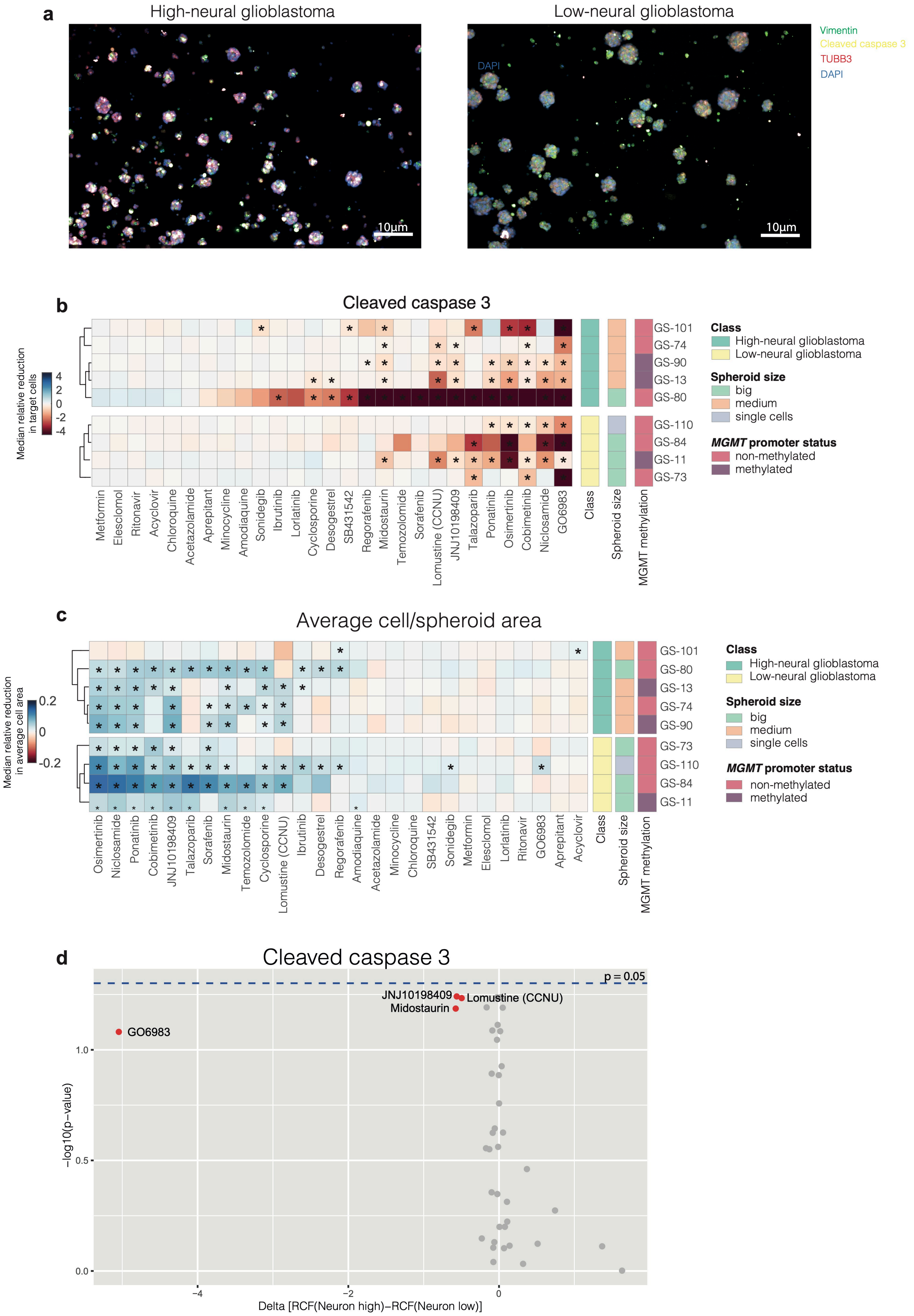

### Supplementary figure 7

**a**

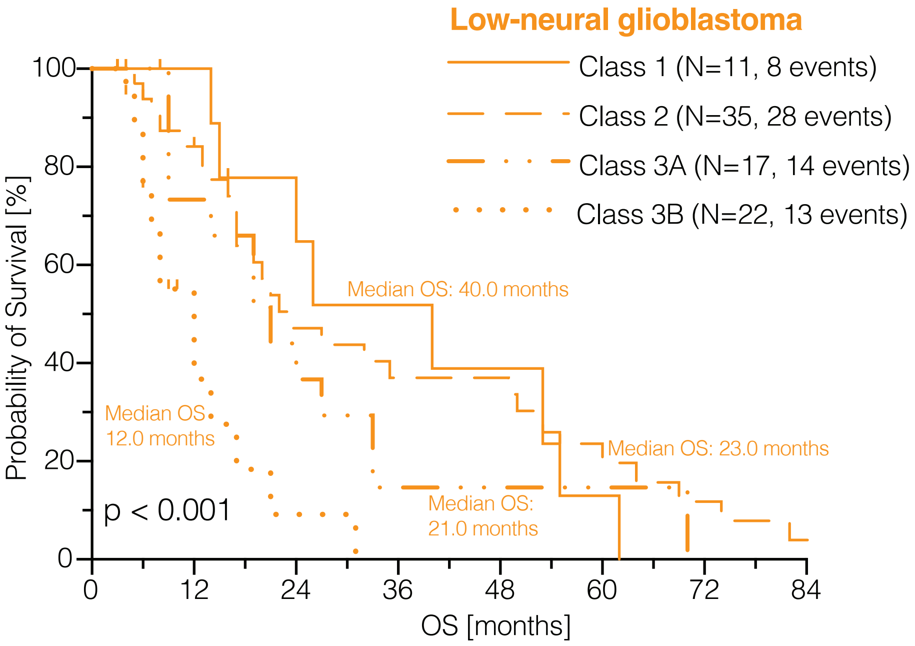


**b**

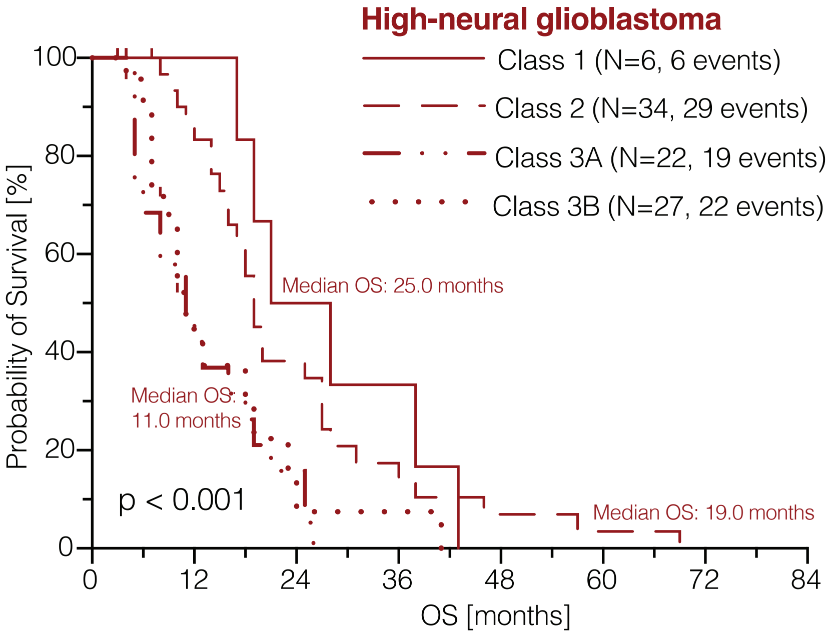

#### Supplementary figure 8

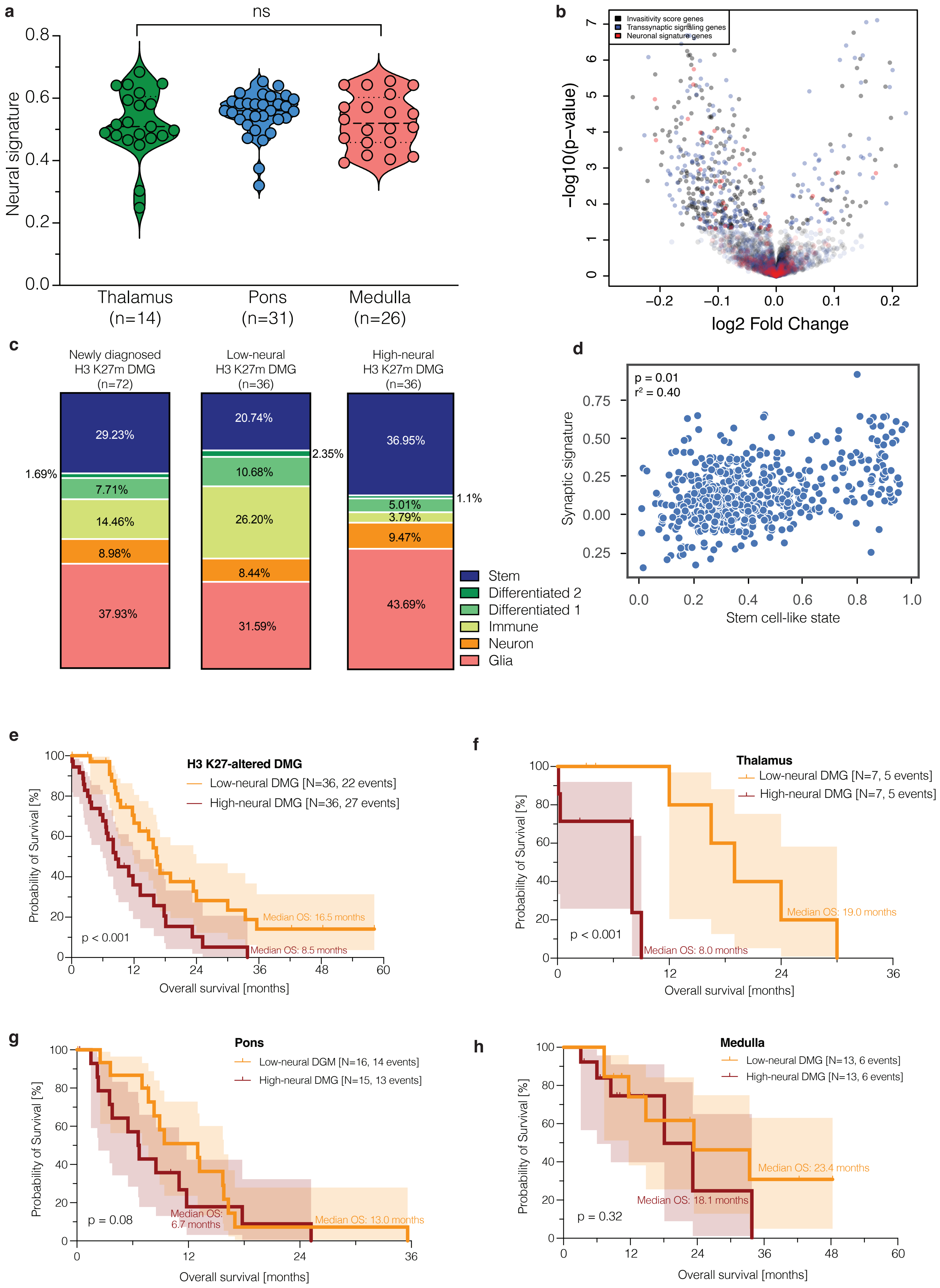
