## Supplementary Table for "Epigenetic neural glioblastoma enhances synaptic integration and predicts therapeutic vulnerability"

**Supplementary Table 1**

| **Characteristic** | **N** | **Low-neural glioblastoma (n=213)** | **High-neural glioblastoma (n=150)** | ***P* value** |
|---|---|---|---|---|
| **Age, mean (SD), years** | 61.4 (10.0) | 60.9 (10.2) | 61.8 (9.7) | 0.41 |
| **Sex, n (%)** | | | | |
| Female | 139 (38.3) | 83 (39.0) | 56 (37.3) | 0.83 |
| Male | 224 (61.7) | 130 (61.0) | 94 (62.7) | |
| **Location, n (%)** | | | | |
| Frontal | 106 (29.2) | 68 (31.9) | 38 (25.3) | 0.19 |
| Parietal | 145 (39.9) | 80 (37.6) | 65 (43.3) | 0.31 |
| Temporal | 141 (38.8) | 71 (33.3) | 70 (46.7) | **0.02** |
| Occipital | 55 (15.2) | 30 (14.1) | 25 (16.7) | 0.55 |
| **Hemisphere, n (%)** | | | | |
| Left | 166 (45.7) | 99 (46.5) | 67 (44.7) | 0.55 |
| Right | 174 (47.9) | 103 (48.4) | 71 (47.3) | |
| Both | 23 (6.3) | 11 (5.2) | 12 (8.0) | |
| **Karnofsky prior surgery, mean (SD), %** | 84.6 (12.4) | 83.0 (12.9) | 86.7 (11.4) | **< 0.01** |
| **Extent of resection, n (%)** | | | | |
| Gross total | 142 (39.1) | 92 (43.2) | 50 (33.3) | 0.08 |
| Near gross total | 99 (27.3) | 59 (27.7) | 40 (26.7) | |
| Partial | 122 (33.6) | 62 (29.1) | 60 (40.0) | |
| ***MGMT* promoter methylation status, n (%)** | | | | |
| Non-methylated | 174 (47.9) | 107 (50.2) | 67 (44.7) | 0.38 |
| Methylated | 189 (52.1) | 106 (49.8) | 83 (55.3) | |
| **Karnofsky prior adjuvant treatment, mean (SD), %** | 85.4 (12.7) | 84.4 (13.0) | 86.7 (12.1) | 0.09 |

### Supplementary Table 2

| Feature | N | Univariate | | Multivariate | |
|---|---|---|---|---|---|
| | | HR (95% CI) | *P* value | HR (95% CI) | *P* value |
| **Age** | 363 | 1.02 (1.00-1.03) | **0.04** | 1.00 (0.98-1.02) | 0.95 |
| **Sex** | | | | | |
| Female | 139 | *Ref.* | | | |
| Male | 224 | 1.11 (0.84-1.47) | 0.46 | | |
| **KPS prior surgery** | 363 | 0.99 (0.98-1.00) | 0.16 | | |
| **Frontal** | | | | | |
| No | 257 | *Ref.* | | | |
| Yes | 106 | 1.08 (0.78-1.24) | 0.53 | | |
| **Temporal** | | | | | |
| No | 222 | *Ref.* | | | |
| Yes | 141 | 0.96 (0.72-1.27) | 0.77 | | |
| **Parietal** | | | | | |
| No | 219 | *Ref.* | | *Ref.* | |
| Yes | 145 | 1.49 (1.11-1.97) | **< 0.01** | 1.26 (0.93-1.71) | 0.46 |
| **Occipital** | | | | | |
| No | 308 | *Ref.* | | *Ref.* | |
| Yes | 55 | 1.46 (1.01-2.11) | **0.04** | 1.04 (0.71-1.52) | 0.86 |
| **Side** | | | | | |
| Left | 166 | Ref. | | | |
| Right | 174 | 1.13 (0.85-1.50) | 0.39 | | |
| Both | 23 | 1.12 (0.59-2.11) | 0.72 | | |
| **Extent of resection** | | | | | |
| GTR | 142 | *Ref.* | | *Ref.* | |
| Near GTR | 99 | 1.38 (0.99-1.94) | **0.04** | 1.14 (0.80-1.63) | 0.46 |
| Partial | 122 | 2.26 (1.61-3.18) | **< 0.01** | 1.84 (1.28-2.64) | **< 0.01** |
| ***MGMT* promoter methylation status** | | | | | |
| Non-methylated | 174 | *Ref.* | | *Ref.* | |
| Methylated | 189 | 0.52 (0.39-0.69) | **< 0.01** | 0.47 (0.35-0.64) | **< 0.01** |
| **KPS prior treatment** | 363 | 0.97 (0.96-0.99) | **< 0.01** | 0.98 (0.96-0.99) | **0.02** |
| **CE volume** | 288 | 0.99 (0.98-1.01) | 0.46 | | |
| **FLAIR volume** | 288 | 1.00 (0.99-1.01) | 0.59 | | |
| **Methylation subclass** | | | | | |
| *RTK I* | 101 | *Ref.* | | | |
| *RTK II* | 147 | 0.95 (0.66-1.36) | 0.77 | | |
| MES | 115 | 0.71 (0.48-1.06) | 0.09 | | |
| **Neural subclass** | | | | | |
| Low | 213 | *Ref.* | | *Ref.* | |
| High | 150 | 1.86 (1.41-2.45) | **< 0.01** | 1.96 (1.45-2.64) | **< 0.01** |

### Supplementary Table 3

| Feature | N | Univariate | | Multivariate | |
|---|---|---|---|---|---|
| | | HR (95% CI) | P value | HR (95% CI) | P value |
| **Age** | 363 | 1.00 (0.99-1.02) | 0.67 | | |
| **Sex** | | | | | |
| Female | 139 | *Ref.* | | | |
| Male | 224 | 0.99 (0.75-1.33) | 0.97 | | |
| **KPS prior surgery** | 363 | 1.01 (0.99-1.02) | 0.22 | | |
| **Frontal** | | | | | |
| No | 257 | *Ref.* | | | |
| Yes | 106 | 1.19 (0.89-1.61) | 0.25 | | |
| **Temporal** | | | | | |
| No | 222 | *Ref.* | | | |
| Yes | 141 | 0.95 (0.71-1.27) | 0.71 | | |
| **Parietal** | | | | | |
| No | 219 | *Ref.* | | | |
| Yes | 145 | 1.17 (0.88-1.56) | 0.28 | | |
| **Occipital** | | | | | |
| No | 308 | *Ref.* | | | |
| Yes | 55 | 1.23 (0.82-1.84) | 0.32 | | |
| **Side** | | | | | |
| Left | 166 | *Ref.* | | *Ref.* | |
| Right | 174 | 1.13 (0.82-1.54) | 0.46 | 1.24 (0.93-1.66) | 0.14 |
| Both | 23 | 1.99 (1.03-3.86) | **0.04** | 1.71 (0.89-3.26) | 0.10 |
| **Extent of resection** | | | | | |
| GTR | 142 | *Ref.* | | *Ref.* | |
| Near GTR | 99 | 1.18 (0.88-1.58) | 0.27 | 1.39 (0.98-1.96) | 0.07 |
| Partial | 122 | 2.29 (1.22-4.29) | **0.01** | 1.73 (1.19-2.49) | **< 0.01** |
| ***MGMT* promoter methylation status** | | | | | |
| Non-methylated | 174 | *Ref.* | | *Ref.* | |
| Methylated | 189 | 0.51 (0.38-0.69) | **< 0.01** | 0.49 (0.37-0.68) | **< 0.01** |
| **KPS prior treatment** | 363 | 0.99 (0.98-1.01) | 0.91 | | |
| **CE volume** | 363 | 1.42 (0.67-3.03) | 0.36 | | |
| **FLAIR volume** | 363 | 1.00 (0.99-1.01) | 0.23 | | |
| **Methylation subclass** | | | | | |
| *RTK I* | 101 | *Ref.* | | | |
| *RTK II* | 147 | 1.11 (0.76-1.62) | 0.60 | | |
| MES | 115 | 1.19 (0.80-1.75) | 0.39 | | |
| **Neural subclass** | | | | | |
| Low | 213 | *Ref.* | | *Ref.* | |
| High | 150 | 1.39 (1.04-1.84) | **0.03** | 1.51 (1.13-2.02) | <0.01 |
